# The structure and global distribution of the endoplasmic reticulum network is actively regulated by lysosomes

**DOI:** 10.1101/2020.01.15.907444

**Authors:** Meng Lu, Francesca W. van Tartwijk, Julie Qiaojin Lin, Wilco Nijenhuis, Pierre Parutto, Marcus Fantham, Charles N. Christensen, Edward Avezov, Christine E. Holt, Alan Tunnacliffe, David Holcman, Lukas C. Kapitein, Gabriele Kaminski Schierle, Clemens F. Kaminski

**Affiliations:** Cambridge Infinitus Research Centre, Department of Chemical Engineering and Biotechnology, University of Cambridge, Cambridge CB3 0AS, United Kingdom; Department of Chemical Engineering and Biotechnology, University of Cambridge, Cambridge CB3 0AS, United Kingdom; UK Dementia Research Institute at the University of Cambridge and Department of Clinical Neurosciences, University of Cambridge, Cambridge CB2 0AH, United Kingdom; Cell Biology, Neurobiology and Biophysics, Department of Biology, Faculty of Science, Utrecht University, Padualaan 8, 3584 CH Utrecht, the Netherlands; Group of Computational Biology and Applied Mathematics, Institut de Biologie de l’École Normale Supérieure-PSL, 46 rue d’Ulm, 75005 Paris, France; Department of Physiology, Development and Neuroscience, University of Cambridge, Cambridge CB2 3DY, United Kingdom; Department of Applied Mathematics and Theoretical Physics, University of Cambridge, Cambridge, CB3 0WA, United Kingdom

## Abstract

The endoplasmic reticulum (ER) comprises morphologically and functionally distinct domains, sheets and interconnected tubules. These domains undergo dynamic reshaping, in response to changes in the cellular environment. However, the mechanisms behind this rapid remodeling within minutes are largely unknown. Here, we report that ER remodeling is actively driven by lysosomes, following lysosome repositioning in response to changes in nutritional status. The anchorage of lysosomes to ER growth tips is critical for ER tubule elongation and connection. We validate this causal link via the chemo- and optogenetically driven re-positioning of lysosomes, which leads to both a redistribution of the ER tubules and its global morphology. Lysosomes sense metabolic change in the cell and regulate ER tubule distribution accordingly. Dysfunction in this mechanism during axonal extension may lead to axonal growth defects. Our results demonstrate a critical role of lysosome-regulated ER dynamics and reshaping in nutrient responses and neuronal development.

## Main text

The structure of the ER is constantly adapted for the particular needs of the cell (*1*): the dynamic transitions between ER sheets and tubules allow it to rapidly respond to the changing cellular environment. A group of ER-shaping proteins have been identified as maintaining ER morphology (*1*), mutations in which are linked to diseases such as hereditary spastic paraplegias (HSPs) (*2*). Whilst these proteins affect global ER topology, they cannot explain the rapid and dramatic changes in ER morphology observable on time scales of minutes in live-cell imaging experiments (*3, 4*). Previous work has shown that ER tubule elongation can be driven by three mechanisms: 1/force generation by motors moving along microtubules (*5*), which can be classified as sliding, 2/coupling to microtubule growth using a tip assembly complex (TAC), and 3/hitchhiking by connecting to other organelles. Whether such reshaping in local domains of the ER tubules could lead to the global reorganization and redistribution of ER remains an open question, and, if this is the case, how is this process regulated? The ER is known to contact other motile organelles, including endosomes, lysosomes, mitochondria, peroxisomes et cetera (*6*). Among these, lysosomes are particularly interesting, as they make a great number of contacts with the ER (*7*) and their positioning is regulated by different nutritional status (*8*). Although ER has been reported to regulate lysosome motions (*9*), it is not clear whether lysosomes can modulate ER reshaping and distribution, for example via coupled motion (*4*). We hypothesized that a causal link exists between lysosome motion and ER redistributing and asked whether this provides a mechanism for ER morphological response to nutritional status, given that lysosomes are known to act as signaling hubs for metabolic sensing (*10*).

We first investigated the correlation of motions between lysosomes and the ER network by rapid live-cell imaging. We visualized ER with GFP-tagged vesicle-associated membrane protein-associated protein A (VAPA) and lysosomes with Cathepsin D-specific SiR-Lysosome dye in COS-7 cells (Fig. 1Ai). Single particle tracking (SPT) of the lysosomes revealed a network of motion tracks resembling the network characteristic of the ER (Fig. 1Aii and Movie S1), featuring path segments, along which lysosomal bidirectional motions take place, and regional nodes, where path segments intersect (Fig. 1Aiii). Compared with lysosomes, early endosomes (labelled with Rab5-GFP) appeared notably more static in COS-7 cells and were significantly less associated with the EGFP-VAPA labelled ER network (Fig. S1 and Movie S2). Tracking lysosomes and the ER network simultaneously using dual-color SPT showed that almost all lysosomes (98%) moved synchronously with local ER domains. This was particularly prominent near elongating tubule tips of the ER (Fig. 1B, Movie S3). Remarkably, almost all lysosome-associated tips (99.4%, 174/175 in 39 cells) eventually formed new connections with the existing ER network, creating three-way junctions (yellow stars in Fig. 1B). In contrast, ER tips without associated lysosomes failed to sustain their elongation (white arrows) and eventually retracted (yellow arrows) (Fig. 1C, Movie S4). These ER tubules normally extended only to about 1 μm, and over half of them (52.5%, 160/305 events in 39 cells) collapsed completely following a retractile phase. Over 90-s periods, ER tubules associated with lysosomes grow significantly longer than their non-associated counterparts (average 6.72±0.269 versus 2.23±0.084 µm, Fig. 1D). In addition, the elongation phase of lysosome-coupled ER tubules persisted longer compared to the lysosome-free cases (9.0 s versus 3.5 s, Fig. 1E). It has previously been reported that 35% of ER tubulation events occur via hitchhiking on late endosomes/lysosomes (*5*), which is consistent with our observation (31%). However, further quantification scaled by tubule length rather than event number shows that 73% of newly generated tubule length was contributed by lysosome-coupled elongation (Fig. 1F). Taken together, these findings suggest that lysosomes play an important role in ER growth tip elongation and the reshaping of local domains within the ER network.

**Fig. 1.**
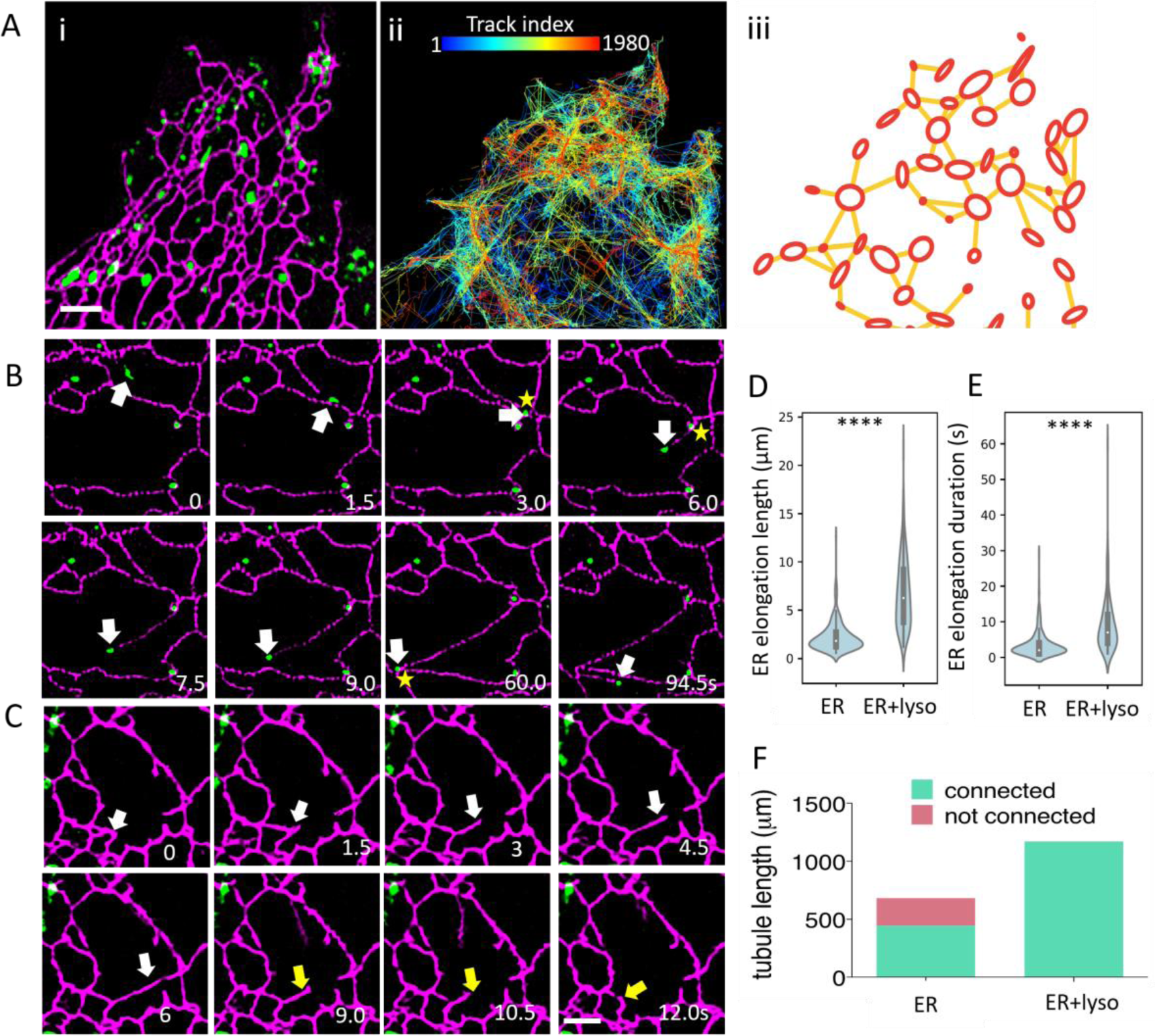
The lysosome-coupled ER network undergoes dynamic morphological changes. **(A) i**: SIM images of ER labelled with EGFP-VAPA (magenta) and lysosomes labeled with SiR dye (green) in a COS-7 cell. **ii**: single-particle tracking (SPT) of lysosome motion over 12.5 mins at 1.5 s/frame, color-coded by track index. **iii**: Track density map extracted from SPT data (see Movie 1 and Methods for details). **(B)** Coupled motion (white arrows) of the growing tip of a newly formed ER tubule (magenta) and associated lysosome (green). The tip forms three-way junctions (yellow stars). See movie S3. **(C)** Time-lapse images of a dynamic ER tubule (arrows), without an associated lysosome. The tubule is unstable and retracts (yellow arrows) after a period of elongation (white arrows). See movie S4. **(D)** ER tubule growth, in a 90 sec interval, for growing tips with or without lysosomes attached. Data are shown as ± SEM. **** = p<0.0001 (student’s t test). **(E)** Maximum duration of the elongation phases of newly formed lysosome-coupled or -free ER tubules. Statistics same as for (**D**). **(F)** Sum of total tubule growth lengths for lysosome-coupled or -free growing ER tips. Light green: tubules making network connections; red: tubules failing to connect. Scale bars represent 2 μm in (**A**), (**B**) and (**C**). s = seconds. For (**D**-**F**), data were collected from 175 events of lysosome-coupled ER motions and 306 events of ER only motions in 39 cells from 3 independent experiments. See table S1.

VAPA has been identified as the main ER-lysosome anchor protein (*11*) and we therefore reasoned that abolishing VAPA-mediated anchoring would lead to compromised ER remodeling. We overexpressed the K87D/M89D (KD/MD) mutant of VAPA, which is unable to bind the FFAT motif of lysosomal membrane proteins and thus compromises the anchoring of the protein to lysosomes based on a previous study (*12*). In this system, we observed frequent detachment of lysosomes from their coupled ER tips during elongation (Fig. 2A, Movie S5). Whilst the lysosome was seen to continue along its trajectory, the ER tubule retracted upon detachment. Remarkably, upon breakage of the link, lysosome speed increased significantly (Fig. 2B), but reduced again once the lysosome reattached to other ER structures, suggesting a resistive force associated with ER growth tip elongation. Silencing VAPA resulted in significant reduction of ER tubules and an extension of the sheet region (Fig. S2), and this ER malformation is consistently observed in the knockdowns of other ER-lysosome anchor proteins, including Protrudin, StAR-related lipid transfer domain containing 3 (STARD3) and oxysterol-binding protein-related protein 1L (ORP1L). Overall, these siRNA treatments of the anchor proteins that are important in maintaining the ER-lysosome contacts suggest that the loss of such contacts lead to a reduced tubular domain in the global ER morphology (Fig. S2).

**Fig. 2.**
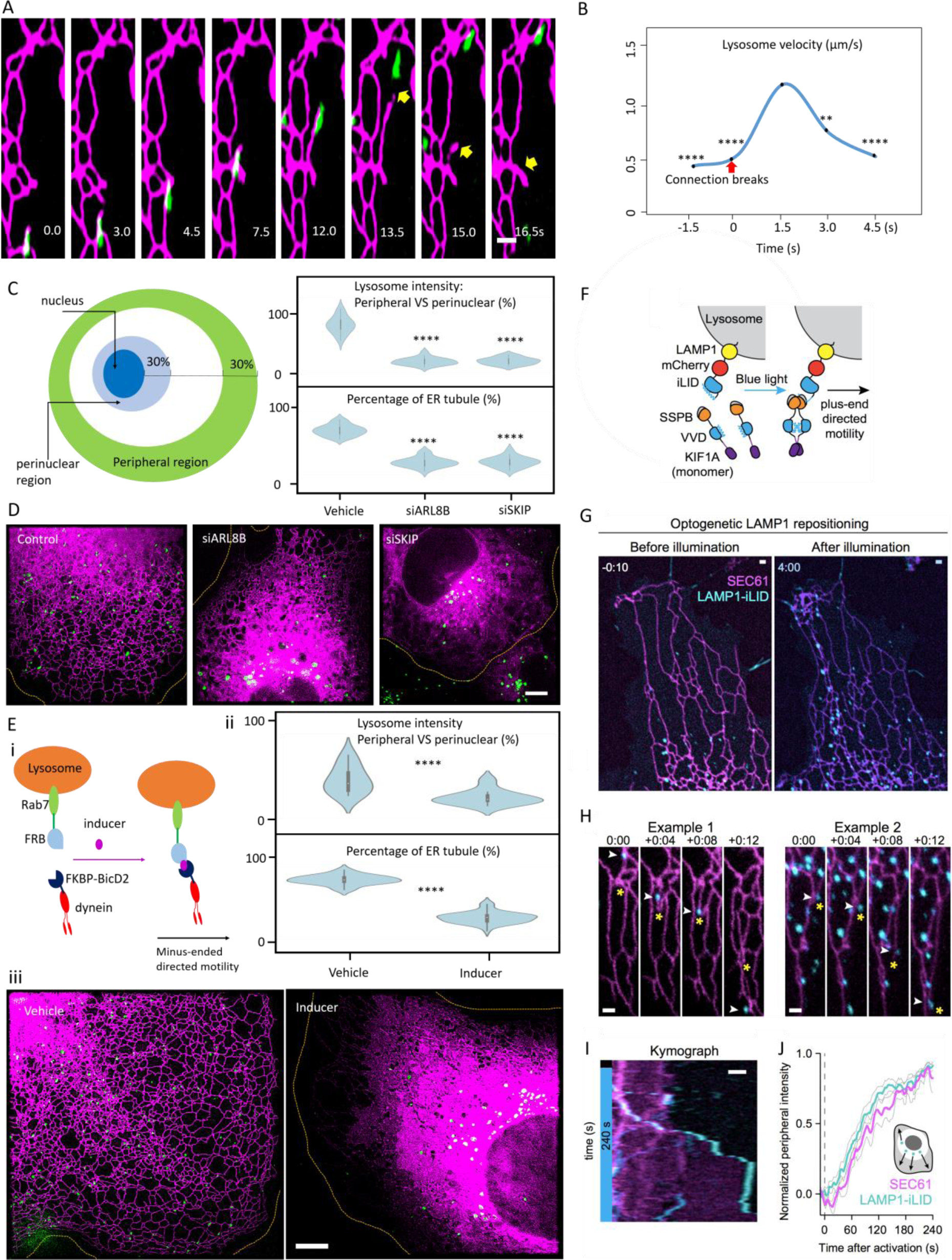
Lysosomes mediate the sustained remodeling of ER networks. **(A)** Representative time-lapse images showing breakage of the connection between the growing tip of a newly-formed ER tubule (magenta) and a lysosome (green) in EGFP-VAPA(KD/MD)-expressing cells. See table S2 for the compromised contact sites upon VAPA(KD/MD) overexpression. See Movie S5. **(B)** Average velocities (black spots) of initially ER-tethered lysosomes that become detached (red arrow) from their associated ER growing tips. Detachment events are ER-lysosome connection breakages in EGFP-VAPA(KD/MD)-expressing cells as in (A). **= p<0.01, **** = p<0.0001 (Tukey’s one-way ANOVA). Velocities of 22 events from three independent experiments were quantified. See table S3. **(C)** Left: Diagram depicting the regions defined as perinuclear and peripheral regions for the following quantification of lysosome distribution, same definition for (**E**). Right: Percentage of the ER comprising tubules upon knockdown of lysosome motion adaptors. Data are shown as ± SEM. **** = p<0.0001 (Tukey’s one-way ANOVA). Data from 20 cells from 3 independent experiments were analyzed for each condition. See table S4. **(D)** Representative images showing the distribution of lysosomes and ER tubules in control cells and in cells treated with siRNAs. **(E) i**: Diagram depicting individual components of the chemogenetic system. **ii**: Quantification of lysosome intensity change and percentage of the ER comprising tubules after 1 hr inducer treatment. Data are shown as ± SEM. **** = p<0.0001 (Student’s t test). For lysosome intensity analysis, *N* = 10, for ER tubule percentage, *N* = 20. **iii**: Representative images showing the distribution of lysosomes and ER tubules in control cells and in cells treated with inducers. **(F)** Optogenetic assay for repositioning of LAMP1. (**G-J**): Live-cell imaging (**G**), representative zoom-ins (**H**), representative kymograph (**I**) and quantification (**J**) of LAMP1-mCherry-iLID and YFP-SEC61B in COS-7 cells expressing opto-kinesin before or during activation. White arrows indicate lysosome pulling ER tubule (yellow asterisk). Quantification shows mean (± S.E.M.) normalized peripheral SEC61 and LAMP1 intensity of 8 cells. Blue box indicates illumination with blue light. Scale bars represent 1 μm in (**A**) (**G**) (**H**) and (**I**), 5 μm in (**D**) and (**E**). s, seconds.

Lysosome positioning is highly regulated by machineries at different nutritional status. ADP ribosylation factor-like GTPase 8B (ARL8B) and SifA-kinesin interacting protein (SKIP), are the key proteins that specifically recruit lysosomes to kinesins for anterograde motions (*13*). We have shown that moving lysosomes are tightly coupled with ER growing tips, pulling ER tubules to move. We therefore hypothesized that compromised anterograde motions of lysosomes can result in the reduction of ER tubules in peripheral regions of the cell. Depletion of ARL8B and SKIP (validated in Fig. S3), which led to perinuclear clustering of lysosomes (Fig2. C and D), strongly reduced the ER tubular domains (Fig2. C and D).

Next, we used the iDimerize Inducible Heterodimer System to examine (*14*) whether the retrograde motion of lysosomes can lead to reduced percentage of tubular network among the whole ER (Fig. 2E). Administration of the inducer led to a recruitment of lysosomes to BicD2 which in turn binds to the dynein complex, thus enabling retrograde transport (Fig. 2Ei). One hour of induction resulted in a significant reduction of lysosome intensity in the peripheral region of the cell and at the same time, the percentage of ER tubule content within the whole network dropped significantly (Fig. 2Eii and iii). We proved in a separate experiment that ER stress alone does not lead to the observed ER morphology changes by administering the ER stress inducer Thapsigargin (Fig. S4).

To further validate the causal link between lysosome repositioning and ER redistribution, we used light-induced heterodimerization to recruit kinesin to LAMP1-labelled lysosomes (*15*) (Fig. 2F). When COS-7 cells co-expressing YFP-ER, KIF1A-VVDfast, an anterograde motor protein, and LAMP1-mCherry-iLlD, engineered LAMP1 with light sensitive protein iLlD, were illuminated with blue light, lysosomes were rapidly redistributed by the recruited kinesin from the perinuclear region to the periphery of the cell (Fig. 2G, see Movie S6 for raw imaging and reconstructed data). Simultaneously, we observed a burst of ER tubule extension towards the periphery in regions where lysosomes were abundant (Fig. 2G). The rapid elongation of ER tubules driven by lysosomes also led to the formation of new network connections (Fig. 2G and H). The kymograph and quantification showed a dramatic and rapid increase in ER fluorescence intensity in cell periphery after blue light illumination, and the growth of ER intensity strictly followed that of lysosome (Fig. 2I and J). The observation of ER tubules growing predominantly in lysosome-enriched regions supports the notion of lysosome-controlled redistribution of the ER network. As lysosomes are the main organelles to sense metabolic stimuli, we asked if this lysosome-guided ER motion is responsive to changing intracellular conditions. We reasoned that intracellular cues that are known to regulate lysosome positioning (Fig. 3A) would result in a redistribution of the ER and transition between tubular and sheet domain. First, we starved COS-7 cells in serum-free culture for 4 hr to induce autophagy, which resulted in the accumulation of lysosomes in perinuclear regions (Fig. 3B). Lysosome movement towards the perinuclear region reduced ER tubules in the cell periphery and a concomitant increase of sheet content in central regions of the cell. In contrast, prolonged serum starvation (24 hr) inhibits autophagy and promotes recycling of lysosomes (*16*), causing their redistribution throughout the whole cytosol. Under this condition, we observed the restoration of the ER tubular network from a previously sheet-dominated state (Fig. 3B). As an example, a dramatic reshaping of the ER is seen to take place in a protrusion forming at a peripheral region of one cell (white box), which is commonly observed under this condition. Motion tracking of lysosomes verified a dominant (>90% among the whole tracks) anterograde motion towards the boxed region (Fig. S5 and Movie S7). However, in the cells treated with siRNA depletion of ARL8B, which led to compromised anterograde motions of lysosomes, ER was not recovered to their native status under prolonged starvation (Fig. 3B). Consistent results were observed in siRNA depletion of SKIP. This provides further evidence that lysosome’s anterograde motion is essential for the ER tubule extension observed above. Quantitative analysis by measuring lysosome intensity and ER tubule percentage showed that accumulation of lysosome in cell center can lead to significant reduction of ER tubules in peripheral regions defined as the regions beyond 70% of the cell radius, and the repositioning of lysosome in cell periphery is accompanied by an extension of ER tubules (Fig. 3C), which is dependent on ARL8B and SKIP for anterograde motions.

**Fig. 3.**
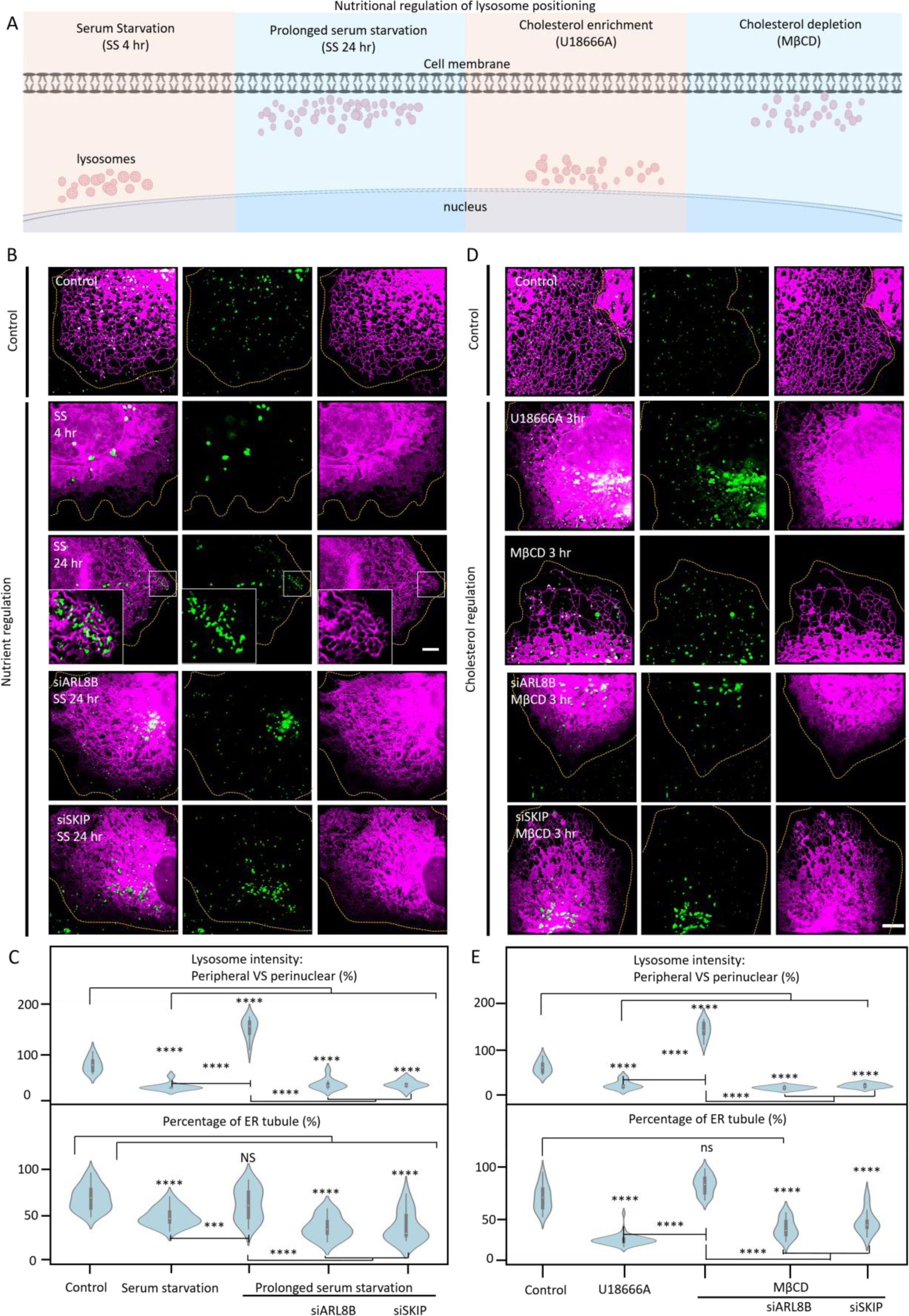
Lysosome-mediated ER reshaping is responsive to nutritional status. **(A)** Schematic representation of the nutritional regulation of lysosome positioning. **(B)** Representative SIM images showing the distribution of lysosomes (green) and ER (magenta) upon disruption of metabolic pathways, same for (**D**). From top to bottom: no treatment control; 4 hr serum starvation (SS, same in the following text); 24 hr serum starvation (prolonged starvation), see Movie S7; siRNA against ARL8B or SKIP in prolonged starvation; **(C)** Quantification of lysosome intensity change and percentage of the ER comprising tubules with different treatments. **(D)** From top to bottom: 3 hr U18666A (10 μM) treatment to block lysosome-to-ER cholesterol transfer; 3 hr MβCD (100 μM) treatment to perturb cholesterol sensing; siRNA against ARL8B or SKIP in 3 hr MβCD (100 μM) treatment. **(E)** Quantification of lysosome intensity change and percentage of the ER comprising tubules with different treatments. Data are shown as ± SEM. * = p<0.05, **** = p<0.0001 (Tukey’s one-way ANOVA). N>=20 cells per condition from 3 independent experiments were analyzed. See table S5. Scale bars: 5 μm.

Lysosomes are known sensors of cholesterol levels in cells, and their positioning is affected by prevailing cholesterol levels (*17*). In order to examine the effects of intracellular cholesterol levels in the context of ER reshaping, we first incubated cells with U18666A, a compound that prevents cholesterol transfer from lysosomes to the ER and leads to lysosome accumulation in perinuclear regions (*18*), which is indeed what we observed upon 3 hr treatment (Fig. 3D). In cells with perinuclear lysosome positioning, the ER also retracted towards the cell center, and showed reduced tubular content and a prevalence sheet structure. In contrast, depletion of cholesterol in COS-7 cells, via administration of Methyl-β-cyclodextrin (MβCD) for 3 hr, prompted a marked redistribution of lysosomes towards the cell periphery and extension of ER tubules in cell periphery (Fig. 3D). However, cells depleted of ARL8B and SKIP failed to extend their ER tubule network in peripheral regions under 3 hr treatment of MβCD, as the lysosomes accumulated in the cell centers (Fig. 3D). Quantitative analysis by measuring lysosome intensity and ER tubule percentage showed that the distribution of ER tubular domain is contingent on lysosome positioning, which is also dependent on ARL8B and SKIP for anterograde motions (Fig. 3E). Further examination by administration of Thapsigargin to COS-7 cells showed that the tubule percentage in the global network is relatively consistent with the vehicle control (Fig. S4). This suggest that the reshaping of ER tubule in Fig. 3B and D is not due to ER stress that may be induced by the metabolic manipulations. In summary, these findings support the notion that lysosome-driven ER reshaping, which leads to the transition between tubular and sheet domain, is responsive to different metabolic activities.

These lysosome-regulated ER redistributions can significantly affect the ER morphology in COS-7 cells, such as the distribution and organization of tubular network and consequently the ratio between tubular and sheet domain. ER-lysosome contacts are also commonly observed in neurons (*19*), in which the ER displays continuous tubular structure in axons (*20*). Whether lysosomes can still drive ER tubule to move and thus maintain ER continuity is not clear, and if so, do they also have any effect on axonal growth? First, to validate the prevalence of lysosome-driven ER motions in a highly polarized system, we studied the cultured axons of *Xenopus laevis* retinal ganglion cell (RGC) neurons, which is a well-established model to study axonal growth (Fig. 4A). Live imaging of ER in RGCs with EGFP-VAPA and lysosomes with LysoTracker showed that 85% (184/217) of lysosomes are coupled with the ER. Consistent with our observation in COS-7 cells, lysosomal pulling of the ER is frequently observed along axons. In Fig. 4Bi, a lysosome (green) is seen to be tethered inside a ring-like structure of a disconnected piece of ER, pulling it along the axon until it reconnects and fuses with an existing ER segment to generate a continuous ER tubule (Movie S8). The lysosome is seen to maintain its anterograde motion after successful ER reconnection. In contrast, in RGCs expressing VAPA (KD/MD), axonal ER ‘repair’ of this type was not seen to take place (Fig. 4Bii and Movie S9). Quantification shows a significant reduction in the reconnection ratio of ER in VAPA (KD/MD)-expressing axons (Fig. 4Biii). To examine whether lysosome-guided ER trafficking is important for healthy axonal growth and development, we transfected RGCs with either EGFP-VAPA or EGFP-VAPA (KD/MD) (Fig. 4C). A quantification of the axon length following overnight growth revealed that in the presence of the VAPA(KD/MD), 24 hr axonal growth was overall compromised compared to the wild type (Fig. 4C and D). The ER–lysosome contacts were maintained all the way along the developing RGC axons, even in the growth cones, as seen in Fig. S6. All these observations suggests that the lysosome-driven ER tubule elongation is a frequent event in axons, which facilitates the ER tubule connections and support axonal growth. The molecular mechanism of how VAPA dependent ER-lysosome coupled motion affects the axonal growth would be interesting for future study.

**Fig. 4.**
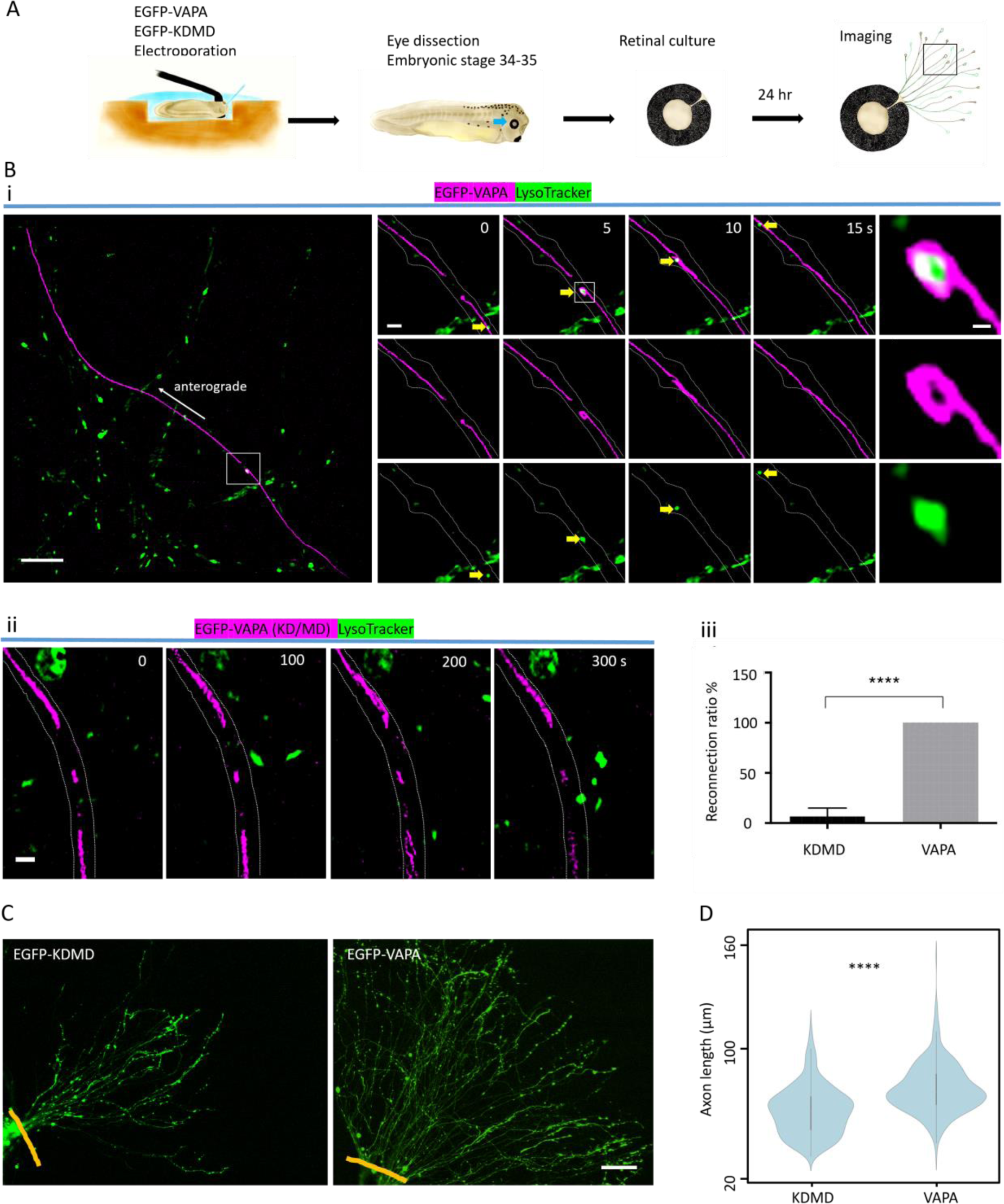
Disruption of ER-lysosome contacts impairs ER continuity in growing axons in RGCs. **(A)** Experimental protocol for *Xenopus* RGC transfection, culture, and imaging. **(B) i**: Representative images showing the co-movement of a moving lysosome (green) and the tip of an ER tubule (magenta) in a *Xenopus* RGC axon (yellow arrows), see Movie S8. Scale bars: 5μm (left), 1μm (middle) and 200 nm (right). **ii**: Representative images showing the persistent ER breakage in an axon, see Movie S9. Scale bar: 1μm. **iii**: Quantification of “repair rate” in ER breakage. Data are shown as ± SEM. **** = p<0.0001 (student’s t test). 221 EGFP-VAPA KD/MD expressing axons from 55 time-lapse movies and 85 EGFP-VAPA expressing axons from 32 time-lapse movies were acquired from three independent experiments. 306 axons and 113 events of ER breakages were quantified. See table S6. **(C)** Representative images of cultured EGFP-VAPA(KD/MD) and EGFP-VAPA expressing RGC axons after 24 h in culture. Yellow lines define the regions where axons start to grow (the starting point of length quantification). Scale bar: 20 μm. **(D)** Quantification of average lengths of EGFP-VAPA(KD/MD)- or EGFP-VAPA-expressing axons after 24 h in culture. Data are shown as ± SEM. **** = p<0.0001 (student’s t test). 434 axons from 12 EGFP-VAPA expressing eyes and 396 axons from 11 EGFP-VAPA KD/MD expressing eyes were quantified from three independent experiments. See table S7.

## Summary and Discussion

Recent advances in live-cell imaging techniques have revealed new details of the highly dynamic nature of the ER structure and its regulation (*3, 4, 5, 21*), posing questions regarding the physiological relevance of this phenomenon. The ER is the main platform for protein and membrane synthesis and quality control of proteins. Lysosomes, on the other hand, are the major sensing and recycling centers, and are actively relocalized in response to cholesterol, nutrient and so on (*10, 22*). Recent studies have revealed that lysosomes can also transport mRNAs (*23*) and act as platforms for protein synthesis (*24*). In this work, we show that movement of ER tubules is actively driven by motile lysosomes, which is facilitated by lysosome-ER anchor proteins. This in turn provides a mechanism for the rapid ER network remodeling required to adapt efficiently to local metabolic changes.

It has been shown that the frequent membrane contact sites enable ER to promote endosome translocation to the cell periphery (*9*), but it is not known whether late endosomes or lysosomes can regulate the global distribution of ER. Our data propose an independent mechanism of regulating ER morphology by lysosome positioning, which is supported by previously reported observations. For example, protrudin, a key anchor protein supporting ER-lysosome contacts, is required to maintain ER sheet-to-tubule balance and the distribution of ER tubules (*25*). Mechanistic study later showed that the anterograde motion of late endosomes promotes protrusion and neurite outgrowth (*9*), which is consistent with our observation of cellular protrusion when lysosome localized to peripheral regions at prolonged serum starvation (Fig. 3B). This suggests a link between lysosome positioning and ER distribution and neurite outgrowth. We further show that the concomitant reorganization of the ER network can result from chemical-, light- and metabolic stimulus-induced lysosome relocation. The dynamic rearrangement of ER tubules is strongly dependent on the motile lysosomes in local domains. These contacts modulate the global ER morphology when lysosomes are regulated to move either retro-/anterogradely at the population level, such as upon cholesterol enrichment and nutrient starvation. Therefore, our results imply that lysosomes sense nutritional changes and regulate ER dynamics and morphology in response. How then ER morphological changes contribute to the maintenance of cellular homeostasis will be an important question for future study.

This lysosome-driven ER redistribution is not only important in ER reshaping, but also facilitate to maintain the continuity of ER in developing axons. A recent study has shown that defects in ER-endosome contacts are associated with HSPs (*26*). In developing RGCs, KD/MD mutants of VAPA, which are incapable of binding to lysosomes, led to persistent ER fragmentation and axonal growth defects, suggesting an important role of lysosomes in maintaining ER continuity during neuronal development. Therefore, this leads to intriguing perspectives in the context of disorders that are primarily linked to ER network defects (*2, 27*). In the future, it will be interesting to study how the lysosome-driven ER dynamics could contribute to the axonal growth and neuronal development.

## Acknowledgements

We thank Dr Ana Isabel Fernández Villegas, Lucia Wunderlich and Wei Li for helping with the cell culture. We thank Dr Edward Ward for helping with the image processing. This research was funded by Infinitus (China) Company Ltd; C. F. K. acknowledges funding from the Wellcome Trust, the UK Medical Research Council (MRC), and Alzheimer Research UK (ARUK). E. A. is supported by the UK Dementia Research Institute which receives its funding from UK DRI Ltd, funded by the UK Medical Research Council, Alzheimer’s Society and Alzheimer’s Research UK. W.N. and L.C.K. acknowledges funding from the Netherlands Organization for Scientific Research (NWO) and the European Research Council (ERC), respectively.

## Supplementary Materials

Fig. S1 to S7

Tables S1 to S8

Captions for Movies S1 to S9

Reference

Movies S1 to S9

## Materials and Methods

### Plasmids

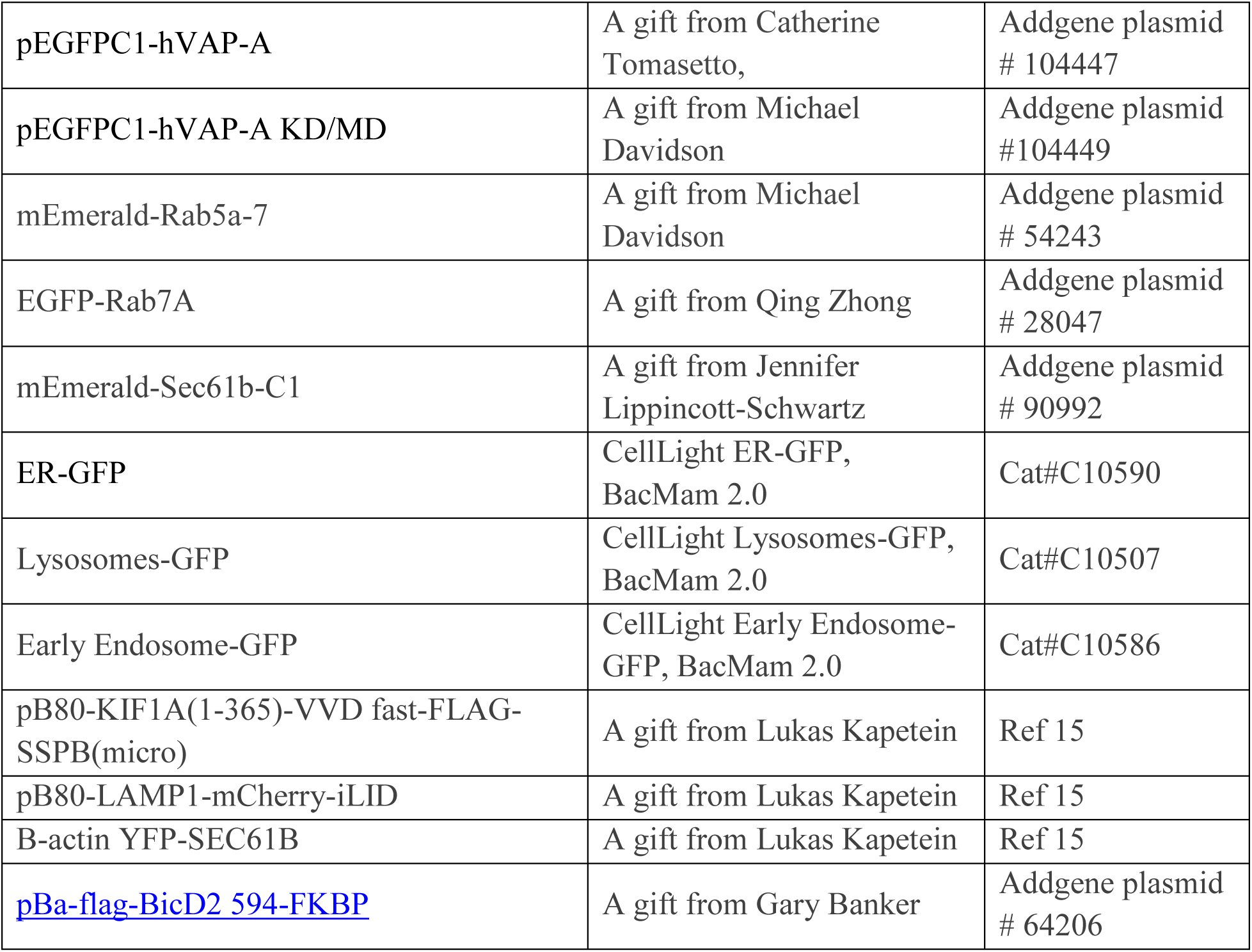

### Antibodies and Chemicals

|  |  |  |
| --- | --- | --- |
| Sir-lysosome | Cytoskeleton, Inc | Cat#CY-SC012 |
| Sir-tubulin | Cytoskeleton, Inc | Cat#CY-SC002 |
| Nocodazole | Sigma-Aldrich | Cat#M1404-10MG |
| Dulbecco's modified eagle's medium (DMEM) | Sigma-Aldrich | Cat#D6546 |
| A/C Heterodimerizer | Takara | Cat#635056 |
| Fetal Bovine Serum (FBS) | Sigma-Aldrich | Cat#10270-106 |
| RIPA buffer | Sigma-Aldrich | Cat#R0278 |
| Opti-MEM™ I Reduced Serum Medium | Invitrogen | Cat#31985062 |
| Anti-VAPA | Abcam | Cat#ab181067 |
| Anti-STARD3 | Abcam | Cat#ab121592 |
| Anti-ORP1L | Abcam | Cat#ab131165 |
| Anti-Protrudin | Proteintech | Cat#12680-1-AP |
| Anti-GAPDH | Sigma-Aldrich | Cat#G8795 |
| Anti-tubulin | Abcam | Cat#ab131205 |
| Anti-PLEKHME | Abcam | Cat#ab91581 |
| Anti-ARL8B | Proteintech | Cat#13049-1-AP |
| Methyl-β-cyclodextrin (MβCD) | Sigma-Aldrich | Cat#332615 |
| U18666A | Sigma-Aldrich | Cat#U3633 |

### Critical Commercial Assays

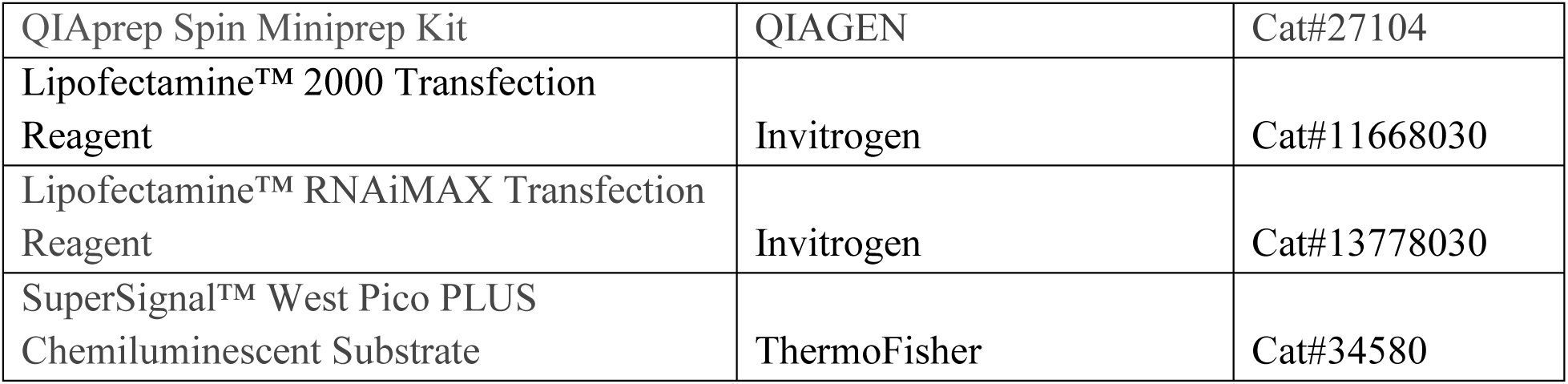

|  |  |  |
| --- | --- | --- |
| QIAprep Spin Miniprep Kit | QIAGEN | Cat#27104 |
| Lipofectamine™ 2000 Transfection Reagent | Invitrogen | Cat#11668030 |
| Lipofectamine™ RNAiMAX Transfection Reagent | Invitrogen | Cat#13778030 |
| SuperSignal™ West Pico PLUS Chemiluminescent Substrate | ThermoFisher | Cat#34580 |

### Experimental Models

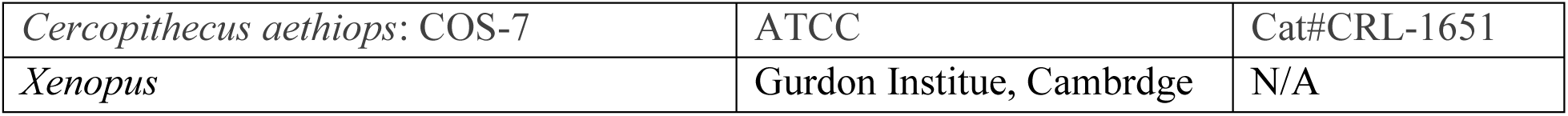

### Oligonucleotides

|  |  |  |
| --- | --- | --- |
| siVAPA | SMARTpool: ON-TARGETplus | Cat#L-021382-00-0005 |
| siProtrudin | SMARTpool: ON-TARGETplus | Cat#L-016349-01-0005 |
| siStard3 | SMARTpool: ON-TARGETplus | Cat#L-017665-00-0005 |
| siOSBIL1A | SMARTpool: ON-TARGETplus | Cat#L-008350-00-0007 |
| MISSION siRNA Universal negative control | Sigma-Aldrich | Cat#SIC001 |
| siPLEKHM2 (SKIP) | SMARTpool: ON-TARGETplus | Cat#L-022168-01-0005 |
| SiARL8B | SMARTpool: ON-TARGETplus | Cat#L-020294-01-0005 |

### Software

|  |  |  |
| --- | --- | --- |
| Fiji | Fiji | https://imagej.net/Fiji |
| MATLAB 2016a | MathWorks | https://uk.mathworks.com/products/matlab.html |
| GraphPad Prism | GraphPad Software | https://www.graphpad.com/scientific-software/prism/ |

### Cell culture

COS-7 cells were purchased from American Type Culture Collection (ATCC). COS-7 cells were grown in T75 or T25 flasks or 6-well plates by incubation at 37°C in a 5% CO_2_ atmosphere. Complete medium for normal cell growth consisted of 90% DMEM, 10% FBS and 1% streptomycin. Cells were kept in logarithmic phase growth and passaged on reaching 70-80% confluence (approximately every 3-4 days). Medium was changed every two or three days. For SIM imaging experiments, COS-7 cells were plated onto Nunc Lab-Tek II Chambered Coverglass (Thermo Scieitific, 12-565-335) to achieve ∼70% confluence prior to transfection.

COS-7 cells were transfected with plasmid constructs as indicated with Lipofectamine 2000 according to the manufacturer’s protocol 24-48 hr before imaging. Cells were stained with SiR-Lysosome at 1 μM 4 hr before imaging. Cells were imaged in a microscope stage top micro-incubator (OKO Lab) with continuous air supply (37°C and 5% CO_2_). Cells were treated with MβCD at100 μM for appropriated time window as indicated. Cells were treated with U18666A at10 μM for appropriated time window as indicated.

#### siRNA transfection and western-blot

VAPA, Protrudin, STARD3, ORP1L were depleted using SMARTpool: ON-TARGETplus, Dharmacon. Negative siRNA control (MISSION siRNA Universal negative control) was purchased from Sigma-Aldrich. COS-7 cells were plated in both glass-bottom petri dishes (for imaging) and 6 well-plate (for western-blot validation). Cells were transfected with 20 nM siRNA oligonucleotides and 20 nM Negative Control siRNA using Lipofectamine(tm) RNAiMax (ThermoFisher Scientific) according to manufacturer’s protocol. After 6 hours of siRNA transfection, the cells were washed and the medium was replaced with complete culture media. 24hr hours after the siRNA transfection, cells were transfected with VAPA-EGFP plasmid DNA using Lipofectamine 2000 (Invitrogen). On the day of imaging, cells were stained with Sir-Lysosome. Cells in glass-petri dishes were imaged 24hr post-DNA transfection.

Cells in 6-well plates were harvested for western-blot validation 72hr post siRNA transfection. Protein concentration was measured using BCA protein assay kit. Immune-blotting was performed by standard SDS-PAGE/Western protocols. Primary <u>antibody concentrations</u> were as follows: VAPA 1:2,0000; Protrudin 1:1000; STARD3 1:500, and ORP1L 1:1000; GAPDH 1:30,000; Tubulin 1:5000. Secondary antibodies (Sigma-Aldrich) was used at 1:3000 for all rabbit antibodies and for all mouse antibodies. Signal was detected with SuperSignal West Pico Chemiluminescent Substrate.

### Structured illumination microscopy (SIM)

SIM imaging was performed using a custom 3-color system built around an Olympus IX71 microscope stage which we have previously described (*28*). Laser wavelengths of 488 nm (iBEAM-SMART-488, Toptica), 561 nm (OBIS 561, Coherent), and 640 nm (MLD 640, Cobolt) were used to excite fluorescence in the samples. The laser beam was expanded to fill the display of a ferroelectric binary SLM (SXGA-3DM, Forth Dimension Displays) to pattern the light with a grating structure. The polarisation of the light was controlled with a Pockels cell (M350-80-01, Conoptics). A 60×/1.2NA water immersion lens (UPLSAPO 60XW, Olympus) focused the structured illumination pattern onto the sample. This lens also captured the samples’ fluorescent emission light before imaging onto an sCMOS camera (C11440, Hamamatsu). The maximum laser intensity at the sample was 20 W/cm^2^. Raw images were acquired with a the HCImage software (Hamamatsu) to record image data to disk and a custom LabView program (freely available upon request) to synchronize the acquisition hardware. Multicolor images were registered via characterising channel displacement using a matrix generated with Tetraspeck beads (Life Technologies) imaged in the same experiment as the cells.

COS-7 cells expressing EGFP-VAPA (ER marker) and stained with SiR-Lysosome (lysosome marker) were imaged by SIM every 1.5 s (including imaging exposure time of both channels) for 1.5 min.

### Reconstruction of the SIM images with LAG SIM

Resolution-enhanced images were reconstructed from the raw SIM data with LAG SIM, a custom plugin for Fiji/ImageJ available in the Fiji Updater. LAG SIM provides an interface to the Java functions provided by fairSIM (*29*). LAG SIM allows users of our custom microscope to quickly iterate through various algorithm input parameters to reproduce SIM images with minimal artefacts; integration with Squirrel (*30*) provides numerical assessment of such reconstruction artefacts.

Furthermore, once appropriate reconstruction parameters have been calculated, LAG SIM provides batch reconstruction of data, so that a folder of multicolor, multi-frame SIM data can be reconstructed overnight with no user input.

### Image analysis algorithms

Further analysis was performed using custom MATLAB algorithms. The displacement of ER network end points, endosomes, and lysosomes was assessed over time with a tracking algorithm. All analysis tools are available from the authors upon request.

### Skeletonization of ER tubules

Skeletonization of images was performed using Fiji (NIH). Images were manually thresholded to reduce the background noise and increase the contrast of ER network. The pre-processed images were imported to Trainable Weka Segmentation (*31*), Fiji, to identify tubules. The trained images were then made binary, and skeletonized by AnalyzeSkeleton (*32*) plugin in Fiji.

### Growing tip and three-way junction analysis

Growing tips and three-way junctions were determined from customer-built algorism. The input to the program is a skeletonized binary image of ER. The skeleton was created in ImageJ by applying a threshold to create a binary image, followed by the built-in ‘skeletonize’ function to reduce the width of ER tubules to 1 pixel. This results in a network graph of the ER, with connections, nodes, and end points. These three features were labelled using MATLAB’s connected components functions. The length of connections was also calculated by fitting splines to the binary tubules and recording the length of the spline. The analysis occurs across a time series of the ER network, allowing results to be drawn about the distribution of tubule length over time, for example. The program ends by saving the labelled image stack as a series of PNG files, as well as several csv files containing statistics describing the network over time.

### ER tubule elongation analysis

ER tubule elongation was counted if the *de novo* ER tubule grew out from the existing ER network. Efficient elongation events are defined as the elongation of tubules over 1 μm and connected with other tubules for over 1 s. Tubular ER generation events were counted if the *de novo* ER tubules branched from the existing ER and extended more than 1 μm. The tubule elongation analysis (length, duration, efficiency and proportions) were quantified from 175 events of lysosome-coupled ER motions and 306 events of ER only motions in 39 cells from 4 independent experiments.

### ER sheet vs tubule ratio

The reconstructed images were imported to Trainable Weka Segmentation (*31*), Fiji, for segmentation of tubule and sheet domains. Channels containing tubules and sheets were then converted to RGB color format. Channels were then split and converted to Mask and measured for the pixel area. For metabolite assay of ER morphology change, 20 images from 3 independent experiments were analyzed for each condition. For siRNA assay of ER morphology change, 20 images from 3 independent experiments were analyzed for each condition.

### Lysosome velocity

To examine the driving force in ER-LY coupled motions, we transfected COS-7 cells with <u>pEGFPC1-hVAP-A</u> KD/MD and measured the velocity change of lysosomes before and after detachment from the coupled ER tips of 23 events from 11 cells from three independent experiments. We measured the displacement of lysosomes between two frames and divided this with time to obtain the lysosome velocity. 23 events from 3 independent experiments were analyzed.

### Single particle tracking

COS-7 cells expressing EGFP-VAPA (ER marker) and stained with SiR-Lysosome (lysosome marker) were imaged by SIM every 1.5 s (including imaging exposure time of SiR-Lysosome channel) for 12.5 min (500 frames). EGFP-VAPA and SiR-Lysosome channels were imaged before and after the recording of lysosome motions, which recorded the morphology of ER. Tracking of lysosome motions was performed by TrackMate (*33*) in Fiji, which plots the trajectories (2094 tracks) of lysosome motions in 500 frames. Data from SPT allowed us to reconstruct the lysosome motion density. The high-density regions were extracted from the trajectories using a procedure derived from (*34*): the number of points falling into square bins of width Δx = 480 nm was counted to construct the density map. From this map, high-density regions, called seeds, were defined as the fraction with 5% higher-density bins. If multiple seeds appeared within a distance of two squares of each other, then only the one with the highest value was kept. For each seed bin, we computed a new density map of size 22 squares, centered on the seed bin center and with bin size Δx/= 200 nm. From this local map, we kept only the ensemble that have a density > 80% of the central bin value. We collected the trajectory points falling into these bins and from which we computed the corresponding ellipse through their covariance matrix (*34*). Finally, when ellipses overlap, we applied an iterative procedure that merge two overlapping ellipses by computing the ellipse based on the ensemble of points falling in each ellipse. The procedure is iterated until there are no more overlaps. From these high-density regions, by considering the displacements connecting different regions, we reconstructed a network explored by the lysosomes, in which a link (yellow) was added when at least one trajectory started in one and entered to the other. Lysosomes appear to traffic between nodes indicated by ellipses.

### Chemogenetics

COS-7 cells expressing mEmerald-Sec61b-C1 (ER marker), pBa-flag-BicD2 594-FKBP and pBa-FRB-3myc-Rab7 (*14*) were imaged by SIM. Lysosomes were stained with SiR-Lysosome as described above. To induce the binding between lysosomes and dyneins, inducer A/C Heterodimerizer was added to the cells at 100 nM for an hour before the imaging.

### Optogenetics

#### Plasmids and cloning

B-actin YFP-SEC61B encodes full length human ER marker SEC61B, fused to EYFP and was a gift from Casper Hoogenraad (Utrecht University, Utrecht, the Netherlands). pB80-KIF1A(1-365)-VVDfast-FLAG-SSPB(micro), encoding opto-kinesin was derived from pB80-KIF1A(1-365)-VVDfast-GFP-SSPB(micro) (15) by replacing GFP with a 2xFLAG-tag (DYKDHDGDYKDHD). pB80-LAMP1-mCherry-iLID was cloned in the mammalian expression vector p*β*actin-16-pl (chicken *β*-actin promoter, referred to hereafter as pB80) (*35*), in which the multiple cloning site was replaced by (*AscI-XbaI-HindIII-NheI-SnaBI-MluI-AgeI*). The full length human lysosome marker LAMP1, without stop codon, was tagged C-terminally with the red fluorescent protein mCherry and the photosensitive heterodimerization module iLID, interspaced by two synthetic 29 amino acid GGGS linkers. The iLID module was derived from pLL7.0-Venus-iLID-Mito (*36*), a gift from Brian Kuhlman (University of North Carolina at Chapel Hill, NC; Addgene plasmid #60413). Full length wildtype human LAMP1, was derived from LAMP1-mGFP (*37*), a gift from Esteban Dell’Angelica (University of California, Los Angeles, CA; Addgene plasmid #34831). All constructs were validated by sequencing of the full ORF.

#### Optogenetic LAMPs repositioning experiments

For live optogenetic LAMP1 repositioning experiments, cells were seeded on 24 mm glass coverslips and transfected with 2 µg plasmid DNA (1:1.3:3 ratio LAMP1:ER:opto-kinesin) and Fugene6 transfection reagent (Promega 1:3) for 20-30 h prior to imaging. Coverslips were mounted in metal rings, immersed in DMEM without phenol red supplemented with 10% FBS, 2 mM L-glutamine and 50 µg/ml penicillin/streptomycin, sealed and maintained at 37°C. Cells were imaged every 2 s for 252 seconds.

Epifluorescence images were acquired using a 40x (Plan Fluor, NA 1.3, Nikon; live-cell imaging) oil-immersion objective on a Nikon Ti inverted microscope equipped with a sample incubator (Tokai-Hit), mercury lamp (Osram), ET-mCherry (49008), and ET 514 nm Laser Bandpass (49905) filter cubes (all Chroma) and a Coolsnap HQ2 CCD camera (Photometrics), controlled with µManager 1.4 software (*38*).

To illuminate the cells during live-cell imaging, we used a Polygon 2000 digital mirror device (DMD) equipped with 405 nm LEDs (Mightex) that exposed the full field of view between imaging frames with an intensity of ∼5 mW cm^-2^. Light exposure was synchronized with camera frames using camera-evoked TTL triggers.

#### Segmentation using deep convolutional neural networks

Segmentation of the tubular networks of the endoplasmic reticulum is done with a convolutional neural network (CNN). The network architecture of choice is a deep residual network inspired by EDSR and RCAN (*39-40*). These models are among a class of residual learning networks (*41-43*) designed for image restoration, specifically single image super-resolution (SR), i.e. image upsampling. The state-of-the-art SR architectures generally do not use downsampling between layers (*35, 37, 38*), but instead make training of deep networks feasible by following the structure of residual networks as first introduced with ResNets (*41*) intended for image classification.

The design idea of residual networks was taken one step further in the EDSR (Enhanced Deep Super-Resolution) model with the proposal of a modified residual building block called ResBlock, which was found made it superior to the previously proposed and more directly adapted ResNet model called SRResNet (*43*).

Another improvement in this class of residual networks was made with RCAN (Residual Channel Attention Network), which augments the ResBlock with two more convolution layers and a global pooling operation. These extra layers are combined with the layers in the standard ResBlock by a modified skip connection that performs multiplication rather addition, which is believed to allow the model to learn to adaptively rescale channel-wise features by considering the interdependencies among channels. RCAN also adds more regular skip connections in a design that is coined Residual in Residual, where a preset number of ResBlocks are considered a group, and a long skip connection is made from the beginning of the group to the end, whereas the skip connection of each ResBlock only passes by a few convolution layers. This allows abundant low-frequency information to be easily passed on, thus enabling the main network to focus on learning to restore high-frequency information.

Choosing the first part of our segmentation model to have an architecture built for restoration ensures that it is capable of handling images with low signal-to-noise ratio as it can learn to perform denoising in these early layers of its network. A neural network model intended for image restoration will by default perform regression in order to output pixel value predictions in the same colour space as the input image. This is achieved during model training by minimising an appropriate loss function, typically the mean squared error defined over the dataset as

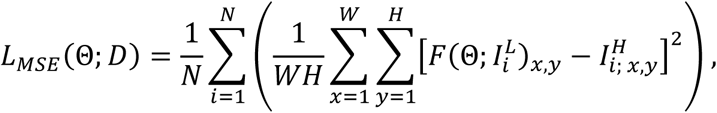

where Θ represents the trainable parameters of the network referred to as *F*(·), while *D* is the training dataset of size *N* written as 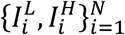 consisting of low-quality input images and high-quality target images with pixel size *H* × *W*.

Rather than having the model perform restoration via regression followed by thresholding by intensity values to produce binary segmentation maps, the model is directly optimised to output segmentation maps by modifying it to perform classification. A common choice of loss function for classification models is the cross-entropy loss given by

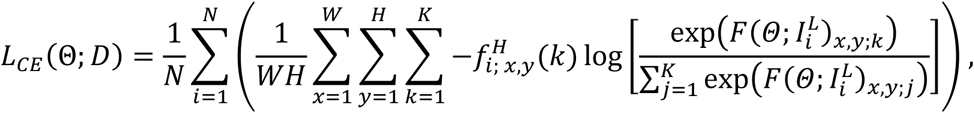

where *k* and *j* are iterators over a total of *K* unique classes, and 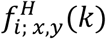 is a function equal to 1 if the target class for the pixel at (*x, y*) of the *i*^th^ image is *k*, and otherwise it is equal to 0. With the model set up for classification, the network *F*(·) now returns scores for each class for a given input image, from which class probabilities are estimated by applying the softmax function, i.e. the normalised exponential inside the log function. The use of the cross-entropy loss over the mean squared error loss during training greatly improves the performance of models with softmax outputs, since the mean squared error tend to lead to saturation and slow learning (*39*), which is why the approach of directly optimising the model to output segmentation maps is preferred. Note that *K* = 2 for the purposes of the binary segmentation used in this work, in which case the innermost summation in *L*_*CE*_(Θ; *D*) over the variable *k* reduces to the addition of two simple terms known as the binary cross-entropy loss.

A proposed network that performs this pixel-wise classification to output segmentation maps is shown on Fig S6. The architecture with a sequence of blocks surrounded by convolutional layers and a long skip connection is the same for both EDSR and RCAN. The definition of the block is different between the two, and for simplicity the EDSR block is shown, although both blocks were used in testing. In general, the RCAN based block was preferred, which. The main difference from the EDSR and RCAN architectures to that of Fig 1 is the replacement of the super-resolution block, an upsampling module, with a module that decodes all the feature channels from previous convolutional layers into class scores. This is done using feature pooling, in which a convolutional layer with kernel size 1−1 reduces the number of feature channels to the number of unique classes in the segmentation map.

Given appropriate training pairs this network can learn to map low signal-to-noise ratio images into clean segmentation maps. Since the network is capable of restoration, it is not necessary to explicitly remove noise (denoise) images prior to inputting them to the model. However, to alleviate the complexity of training the network, raw images were first denoised using the denoising method ND-SAFIR (*45*) which includes a noise parameter estimation of Poisson-Gaussian noise that is typical for optical microscopy.

As for preparing the training data, a very crude segmentation could be performed using grayscale pixel intensity thresholding after applying this denoising method to raw images. A few of these segmentation maps were then manually cleaned and finalized by drawing in a raster graphics editor. These partially hand-drawn segmentation maps then served as targets, i.e. ground truths, in the supervised training of the segmentation network.

To make it feasible for the network to learn to segment the ER images from the relatively small training dataset, a few means of data augmentation were used. Firstly, each segmentation training pair was randomly cropped many times, which shifts the structures from frame to frame and brings the image size down to a manageable size for a graphics card (256−256 pixels). This provides a few hundred subimages of different regions of the segmentation examples. Those subimages were then randomly flipped (horizontally and vertically) and rotated (by 90 or 180 degrees) to obtain more training data. Other ways to augment data could have been to randomly change brightness or synthetically add noise to images, but this was not found to be necessary.

Training was done in batches of 5 images (each being 256−256 pixels) with a learning rate of 0.001 using the Adam optimisation method for a total of 40 epochs. The learning rate was halved after every 10 epochs. The network was conFigd to have 4 blocks, of the RCAB, which amounts to 40 convolution layers each having 64 filters with a 3−3 kernel size, constituting a total of about 1.3 million trainable parameters. The trained network outputs binary segmentation maps that can then easily be skeletonised by a standard thinning algorithm (*46*).

The implementation has been made with the machine learning library Pytorch. The code as well as training and test data for segmenting endoplasmic reticulum images is freely available at https://github.com/charlesnchr/ERNet.

#### Targeted eye electroporation of *Xenopus*

Targeted eye electroporation can be performed as previously described (*47-48*). To anesthetize embryos during electroporation, 0.4 mg/ml tricaine methanesulfonate (MS-222) (Sigma) in 1x MBS, pH 7.5, is used. An anesthetized embryo is positioned along the longitudinal channel of a “†” shape electroporation Sylgard chamber, with its head positioned at the cross of the longitudinal and transverse channels. A pair of flat-ended platinum electrodes (Sigma) is held in place by a manual micromanipulator (World Precision Instruments) at the ends of the transverse channel. A glass capillary with a fine tip containing plasmid solution is inserted into the eye primordium of stage 26-30 embryos to inject 8−5-8 nl doses of 1 µg/µl of pEGFPC1-hVAP-A (Addgene #104447) or pEGFPC1-hVAP-A KD/MD (Addgene #104449) plasmid driven by an air-pressured injector, such as a Picospritzer (Parker Hannifin). Immediately following the plasmid injection, 8 electric pulses of 50 ms duration at 1000 ms intervals are delivered at 18 V by a square wave generator, such as the TSS20 OVODYNE electroporator (Intracel). The embryos are recovered and raised in 0.1x MBS until they reach stage 32-35 as required for retinal cultures.

#### *Xenopus* retinal culture and Imaging

Glass bottom dishes (MatTek) are pre-treated with 5 M KOH (Sigma) for 1 hour at room temperature to minimize background fluorescence, followed by 5-8 rinses with deionized water (Sigma), and are then left to dry in a hood (all culturing procedures should be performed in a laminar flow hood or a microbiological safety cabinet). They are next coated with 10 µg/ml poly-L-lysine (Sigma) overnight at room temperature. Excess poly-L-lysine is discarded from the dishes, which are then rinsed three times with double-distilled water and left in the hood until dry. Subsequently, the dishes are coated with 10 µg/ml laminin (Sigma) in L15 (GIBCO) for 1-3 hours. Finally, the laminin solution is replaced with culture medium (60% (v/v) of L15, 1% (v/v) Antibiotic–antimycotic (100x), in double-distilled water, pH 7.6-7.8, sterilized with 0.22µm pore-size filters), in which dissected eyes can be placed.

Electroporated embryos at stage 32-35 are first screened for EGFP fluorescence to check for successful electroporation. Embryos with fluorescent eyes are then rinsed three times in the embryo wash solution (0.1x MBS with 1% (v/v) Antibiotic–antimycotic (100x) (Thermo Fisher Scientific), pH 7.5, sterilized with 0.22µm pore-size filters) and anesthetized in MS-222 solution (0.04% (w/v) MS-222 and 1% (v/v) Antibiotic–antimycotic (100×) in 1x MBS, pH 7.5, filtered with 0.22µm filters). After transfer of the embryos to the Sylgard dish, the electroporated eye primordia are dissected out and washed three times in culture medium. Finally, the dissected eye primordia are placed in the center of the dish, and the cultures are incubated at room temperature overnight to allow axon extension.

#### Wide-field imaging of *Xenopus* axons and analysis

24 hr after culturing, fluorescent RGC axons are identified and imaged under an Olympus IX81 inverted microscope with a 20X 0.45 NA air objective at room temperature. As the axons extended beyond the field of view, multiple positions of axons projecting from the same eye are imaged to cover all the axons.

Images of *Xenupus* RGC axons are processed and analyzed using Fiji (NIH). Images at multi-positions are mapped and integrated. EGFP-labeled axons are manually traced by “Freehand line” from the distal tips to the proximal segments until reaching the explants. The axon length is quantified by “Measure-Analyze” in Fiji. Three independent experiments were performed, in which 446 axons from 12 EGFP-hVAPA expressing eyes and 396 axons from 11 EGFP-hVAPA KD/MD expressing eyes were quantified.

#### SIM imaging of *Xenopus* RGC axons

To visualize lysosomes, retinal cultures are incubated with 50 nM LysoTracker Deep Red (ThermoFisher Scientific) for 30 minutes at room temperature and rinsed 3 times with culture medium. Time-lapse recordings of ER and lysosomes are taken at using a custom 3-color system built around an Olympus IX71 microscope stage which have been described above. Laser wavelengths of 488 nm (iBEAM-SMART-488, Toptica) and 640 nm (MLD 640, Cobolt) were used to excite fluorescence in the samples. Multicolor images were registered via characterising channel displacement using a matrix generated with Tetraspeck beads (Life Technologies) imaged in the same experiment as the cells.

Samples were imaged at 3 or 5 s per frame for 60 frames immediately following the LysoTracker staining. Resolution-enhanced images were reconstructed from the raw SIM data with LAG SIM as described above. The axon outlines were drawn via increasing the brightness of fluorescence from LysoTracker staining in the analysis step. 221 EGFP-hVAPA KD/MD expressing axons from 55 time-lapse images and 85 EGFP-hVAPA expressing axons from 32 time-lapse images were acquired from three independent experiments.

#### Data visualization

Videos of time-lapse imaging and analysis were performed using Fiji (NIH).

#### Statistical Analysis

Statistical significance between two values was determined using a two-tailed, unpaired Student *t*-test (Graphpad Prism). Statistical analysis of three or more values was performed by one-way analysis of variance with Tukey’s *post hoc* test (Graphpad Prism). All data are presented as the mean ± standard error of the mean; \*\*\**P*<0.001,\*\**P*<0.01.

Statistical parameters including the exact value of n, the mean, median, <u>dispersion</u> and precision measures (mean ± SEM) and statistical significance are reported in the Figs and Fig Legends. Data is judged to be statistically significant when p < 0.05 by two-tailed Student’s t test. In Figs, asterisks denote statistical significance as calculated by Student’s t test (*, p < 0.05; **, p < 0.005; ***, p < 0.0005). Statistical analysis was performed using both Origin and Excel unpaired t test.

