## Supplementary Materials for "The structure and global distribution of the endoplasmic reticulum network is actively regulated by lysosomes"

**This file includes:**

Fig. S1 to S7
Tables S1 to S8
Captions for Movies S1 to S9
Reference

**Other supplementary material for this manuscript includes the following:**

Movies S1 to S9

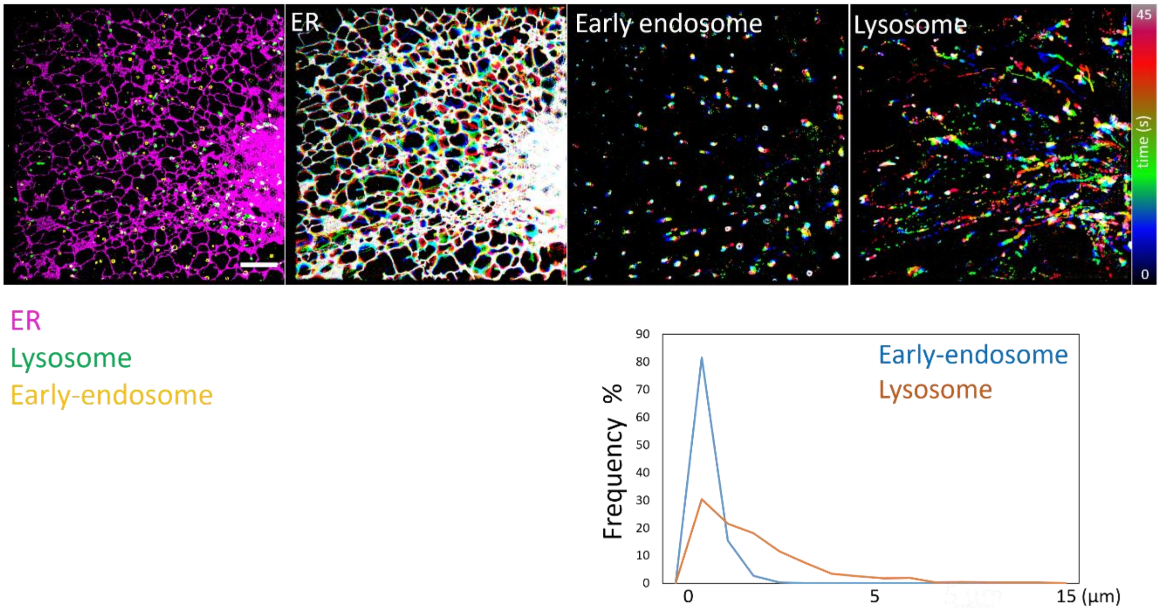


**Fig. S1. Comparison of organelle dynamics between early endosome and lysosome.**
First left: SIM image from a time-lapse recording of a COS-7 cell expressing Rab5-EGFP (yellow) and mApple-Sec61b-C1 (magenta), stained with SiR-lysosome (green). From second left to right: images of ER, early endosome, and lysosome, all color coded to represent time. (**B**) Histograms of net displacement of early endosomes and lysosomes during a recording time of 90 s. 3025 early endosomes and 1000 lysosomes were analysed from three different experiments. Scale bar: 5 µm. (see Movie S2 and table S8).

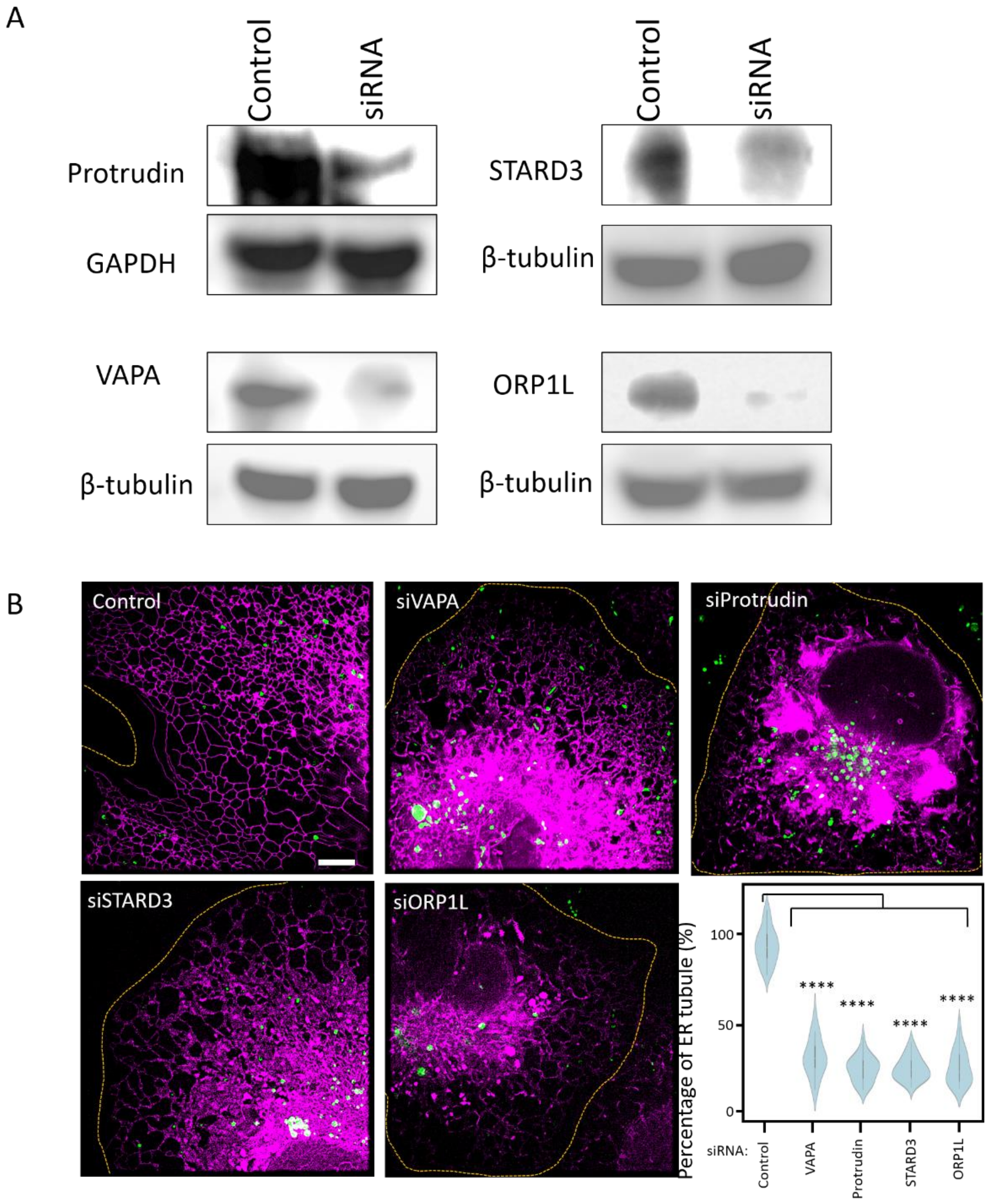


**Fig. S2. Depletion of ER-lysosome anchor proteins lead to ER malformation.**
(**A**) Western blots to confirm depletion of VAPA, Protrudin, STARD3, and ORP1L. *N*=2
(**B**) Spatial distribution of ER labeled by Sec16β-EGFP (magenta) and lysosomes labeled by SiR-Lysosome (green) in COS-7 cells with different ER-lysosome anchoring complex genes knocked down by siRNA. Bottom right: Percentage of the ER comprising tubules upon knockdown of different ER-lysosome anchoring genes. Data are shown as ± SEM. **** = $p<0.0001$ (Tukey's one-way ANOVA). Data from 20 images from 3 independent experiments were analyzed for each condition. See table S4.

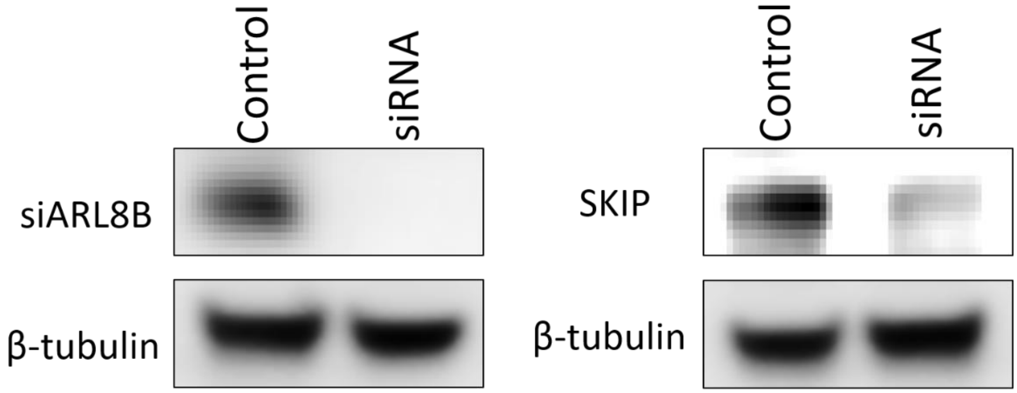


**Fig. S3. Western blots to confirm depletion of ARL8B and SKIP.**

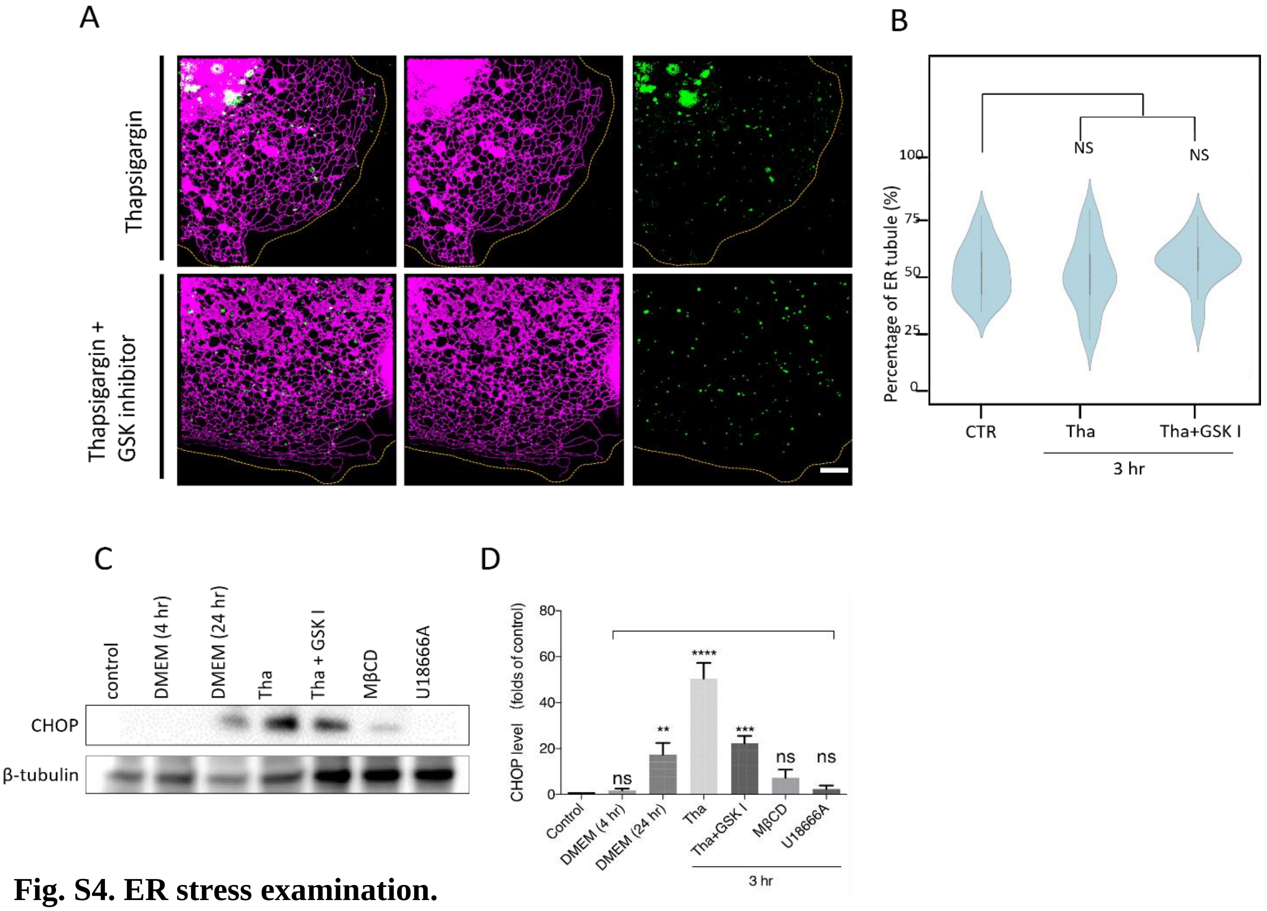


**Fig. S4. ER stress examination.**

(**A**) Representative COS-7 cells expressing Sec16β-EGFP (magenta) and stained with Sir-Lysosome (green), after 3 h of Thapsigargin treatment (500 nM) or Thapsigargin (500 nM) plus GSK inhibitor (5 μM). (**B**) Quantification of ER tubule percentage under the two treatment. (**C**)

Immunoblot analysis with antibodies against CHOP revealed elevation of ER stress in Thapsigargin treated COS-7 cells. (**D**) Quantification of the CHOP levels in immunoblots (*N*=3).

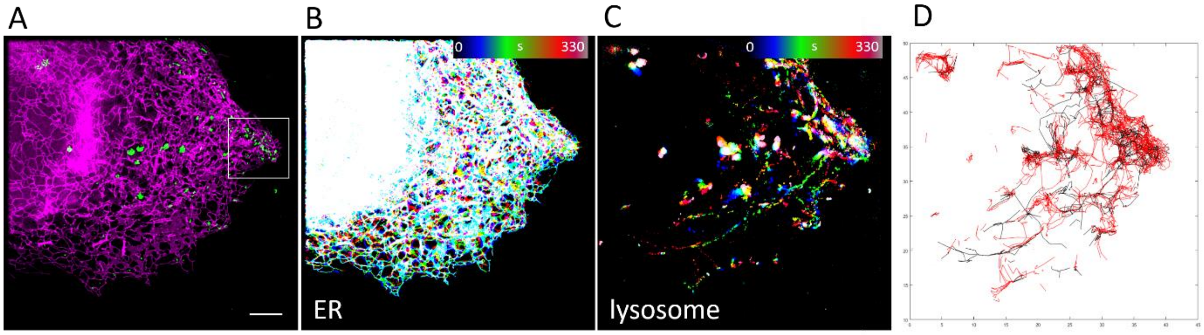


**Fig. S5. Motion tracking of lysosomes in prolonged serum starvation.**
(**A**) Representative COS-7 cells expressing Sec16β-EGFP (magenta) and stained with Sir-Lysosome (green), after 24 h of DMEM treatment. Images are color-coded to represent time and show, (**B**), ER and ,(**C**), lysosome motion. (**D**) Lysosome tracks, color coded according to direction of motion. Red: lysosome move towards the protrusion highlighted by the white box. Black: lysosome tracks that move away from the protrusion. Units of x- and y-axes: µm. Scale bar: 5 µm. (See Movie S7)

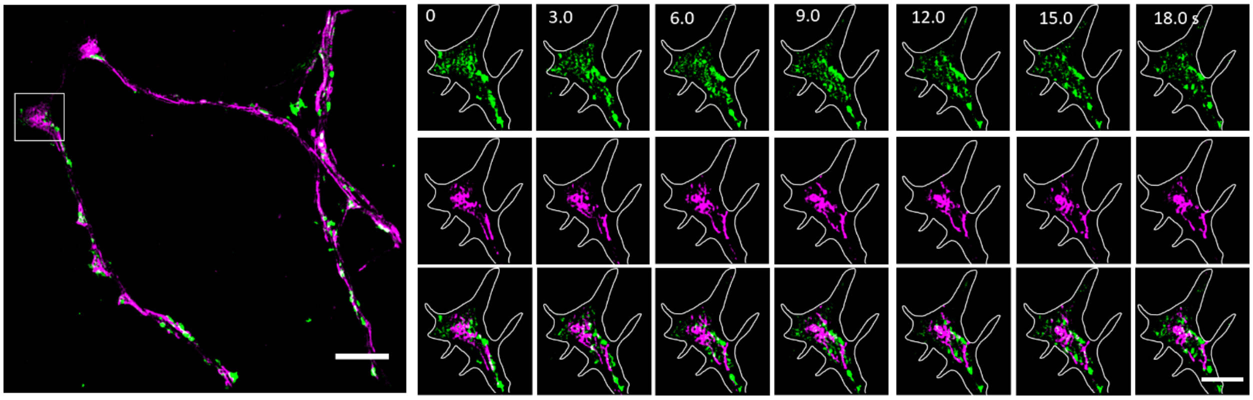


**Fig. S6. Colocalization of lysosomes and ER in a *Xenopus laevis* growth cone.**
Representative images showing stable contacts of lysosomes and ER in an RGC growth cone. During the whole recording of 18 s, lysosomes remained attached to the ER. Scale bar: 5 µm (left) and 2 µm (right).

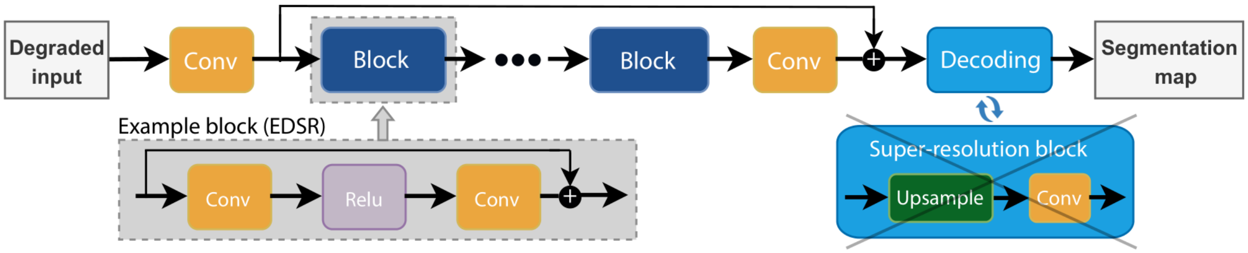


**Fig S7**. Architecture of the residual CNN used for segmentation. The overall structure follows that of EDSR and RCAN, except for the replacement of the super-resolution block with a decoder module that reduces the number of feature channels to the number of unique classes in the segmentation map using a convolutional layer with a corresponding number of output channels and a kernel size of 1x1. This operation is sometimes referred to as feature pooling.

**Table S1.**

Quantification of ER elongation events in Fig. 1D and E. Three independent experiments were performed to measure the ER tubule elongation length, duration and efficiency.

Fig. 1D

| **Condition** | ***N* (cells)** | ***N* (events)** | **Elongation length ± SD (μm)** |
|---|---|---|---|
| ER | 39 | 305 | 2.23±0.084 |
| ER+LY | 39 | 175 | 6.72±0.269 |

Fig. 1E

| **Condition** | ***N* (cells)** | ***N* (events)** | **Elongation duration ± SD (s)** |
|---|---|---|---|
| ER | 39 | 305 | 3.54 |
| ER+LY | 39 | 175 | 8.92 |

Fig. 1F

| **Condition** | ***N* (cells)** | ***N* (events)** | **Successful elongation length (μm)** | **Unsuccessful elongation length (μm)** |
|---|---|---|---|---|
| ER | 39 | 305 | 442.92 | 237.18 |
| ER+LY | 39 | 175 | 1169.73 | 1.77 |
| Total | 39 | | 1612.65 | 238.95 |

| **Experiment 1** | | **ER** | **ER+LY** |
|---|---|---|---|
| *N* (cells) | | 23 | 23 |
| *N* (events) | | 165 | 105 |
| Successful elongation length (μm) | A | 246.87 | 719.62 |
| Unsuccessful elongation length (μm) | B | 130.32 | 1.76 |
| Elongation efficiency (%) | A/(A+B) | 65.44 | 99.75 |
| Elongation duration (s) | | 530 | 931 |

| **Experiment 2** | | **ER** | **ER+LY** |
|---|---|---|---|
| *N* (cells) | | 6 | 6 |
| *N* (events) | | 59 | 27 |
| Successful elongation length (μm) | A | 80.705 | 167.195 |
| Unsuccessful elongation length (μm) | B | 41.127 | 0 |
| Elongation efficiency (%) | A/(A+B) | 66.24 | 100 |
| Elongation duration (s) | | 83.5 | 166 |

| **Experiment 3** | | **ER** | **ER+LY** |
|---|---|---|---|
| *N* (cells) | | 10 | 10 |
| *N* (events) | | 81 | 43 |
| Successful elongation length (μm) | A | 115.341 | 282.911 |
| Unsuccessful elongation length (μm) | B | 65.721 | 0 |
| Elongation efficiency (%) | A/(A+B) | 63.70 | 100 |
| Elongation duration (s) | | 259.5 | 445.5 |

**Table S2.**

Quantification of ER-LY coupled motion in either EGFP-hVAPA KD/MD or EGFP-hVAPA expressing COS-7 cells. Expression of KD/MD mutant of VAPA resulted a significant proportion of lysosomes detached from ER during motions (66%). As noted in the following table, the proportion of lysosomes contacted with ER also decreased from 97.7% (VAPA) to 75.5% (KD/MD). Fig. D-F

| | **VAPA** | **KD/MD** |
|---|---|---|
| Total lysosome No. | 710 | 634 |
| Lysosomes connected with ER | 694 | 479 |
| Lysosomes coupled with ER in motions | 70 | 56 |
| Detached lysosomes from coupled motions | 0 | 37 |
| *N* (cells) | 11 | 10 |
| *N* (repeats) | 2 | 2 |

**Table S3.**

Quantification of lysosome velocity in the events of ER-LY detachment. Three independent experiments were performed to measure the velocities of lysosomes at five continuous time points. Fig. 2B

| **Time point (s)** | **-1.5** | **0** | **1.5** | **3** | **4.5** |
|---|---|---|---|---|---|
| average velocity (μm/s) | 0.390712 | 0.485424 | 1.047333 | 0.65 | 0.46703 |
| *N* (events) | 22 | 22 | 22 | 22 | 22 |
| *N* (cells) | 15 | 15 | 15 | 15 | 15 |
| *N* (repeats) | 3 | 3 | 3 | 3 | 3 |

**Table S4.**

Quantification of tubule domain among the whole ER in two independent experiments of different gene knock-downs by siRNA. Fig. 2C and S2.

| **Treatment** | **CTR (control)** | **siVAPA** | **siProtrudin** | **siSTARD3** | **siORP1L** | **siARL8B** | **siSKIP** |
|---|---|---|---|---|---|---|---|
| Percentage of ER tubules (%) | 80.496315 | 19.80632 | 13.3578 | 13.5024 | 13.98494 | 24.32851 | 25.12999 |
| N (cells) | 40 | 20 | 28 | 29 | 20 | 20 | 20 |
| N (repeats) | 2 | 2 | 2 | 2 | 2 | 2 | 2 |

**Table S5.**
Quantification of tubule domain among the whole ER in in two independent experiments of different metabolic treatments.
Fig. 3C and E.

| Treatment | CONTROL | DMEM 4hr | DMEM 24 hr | siARL8B DMEM 24 hr | siSKIP DMEM 24 hr | U18666A | MβCD | siARL8B MβCD | siSKIP MβCD |
|---|---|---|---|---|---|---|---|---|---|
| Percentage of ER tubules (%) | 61.83 | 38.31 | 54.45 | 24.93 | 25.91 | 9.27 | 78.15 | 23.96 | 31.70 |
| N (cells) | 41 | 29 | 27 | 15 | 15 | 47 | 34 | 15 | 16 |
| N (repeats) | 2 | 2 | 2 | 2 | 2 | 2 | 2 | 2 | 2 |

**Table S6.**
Quantification of ER breakages and connections in either EGFP-hVAPA KD/MD or EGFP-hVAPA expressing RGC axons. Four independent experiments were performed to measure the number of ER tubule breakages and connections, either via ER only or lysosome-driven elongation.
Fig. 4Biii

***KD/MD***

| Experiment | *N* (axons) | ER breakage gaps | Gap connection | Lysosome-driven connection |
|---|---|---|---|---|
| 1 | 92 | 51 | 2 | 0 |
| 2 | 34 | 8 | 0 | 0 |
| 3 | 89 | 44 | 7 | 2 |
| 4 | 6 | 0 | 0 | 0 |
| Total | 221 | 103 | 9 | 2 |

***VAPA***

| Experiment | *N* (axons) | ER breakage gaps | Gap connection | Lysosome-driven connection |
|---|---|---|---|---|
| 1 | 40 | 3 | 3 | 2 |
| 2 | 17 | 0 | 0 | 0 |
| 3 | 18 | 4 | 4 | 3 |
| 4 | 10 | 3 | 3 | 1 |
| Total | 85 | 10 | 10 | 6 |

**Table S7.**

Quantification of length of either EGFP-hVAPA KD/MD or EGFP-hVAPA expressing RGC axons after overnight culture. Three independent experiments were performed.

Fig. 4D

| | KD/MD | | VAPA | |
|---|---|---|---|---|
| Experiment | N (axons) | Length ± SD (μm) | N (axons) | Length ± SD (μm) |
| 1 | 199 | 101.30 | 187 | 130.23 |
| 2 | 207 | 91.00 | 167 | 136.29 |
| 3 | 28 | 86.80 | 42 | 119.46 |

**Table S8.**

Comparison of the displacements between early endosomes and lysosomes. Three independent experiments of time-lapse Fast-SIM imaging were performed to measure the displacements of each motion tracks via TrackMate, Fiji.

Fig. S1

| | Early endosome | | Lysosome (Late endosome) | |
|---|---|---|---|---|
| Experiment | N (tracks) | Displacement ± SD (μm) | N (tracks) | Displacement ± SD (μm) |
| 1 | 1039 | 0.57 | 328 | 2.46 |
| 2 | 1036 | 0.43 | 273 | 3.02 |
| 3 | 947 | 0.50 | 399 | 2.28 |

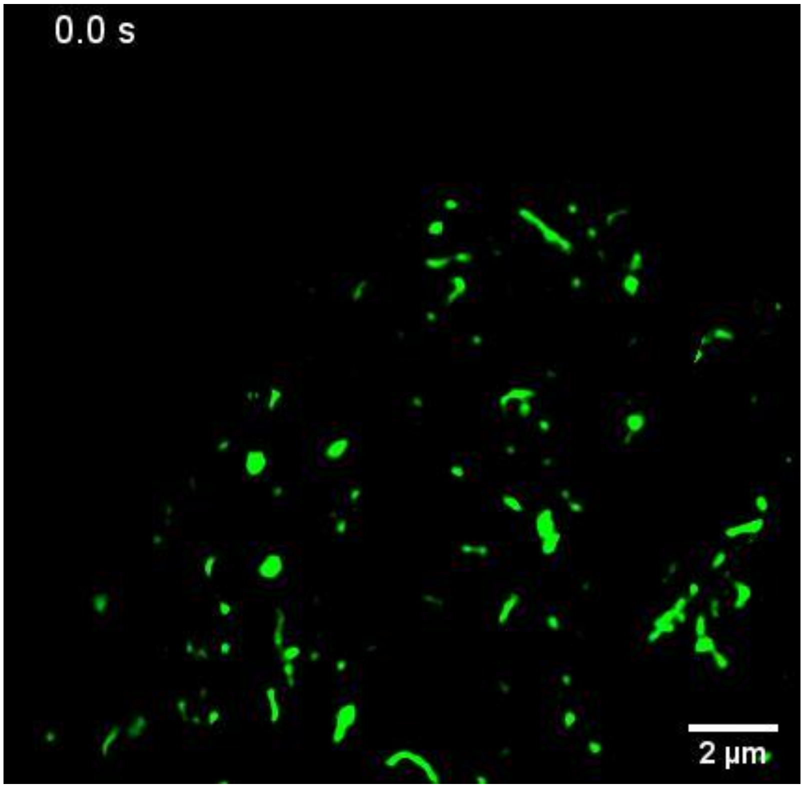


**Movie S1.**

**Live cell imaging by SIM demonstrates the dynamic motions of lysosomes.** A representative 100 frames of whole recording. COS-7 cells were stained by SiR-Lysosome and a region was imaged over 12.5 mins at 1.5 s/frame, from which Trackmate was used to extract lysosome positions at each time point for further analysis. (see **Fig. 1Aii** and **iii**).

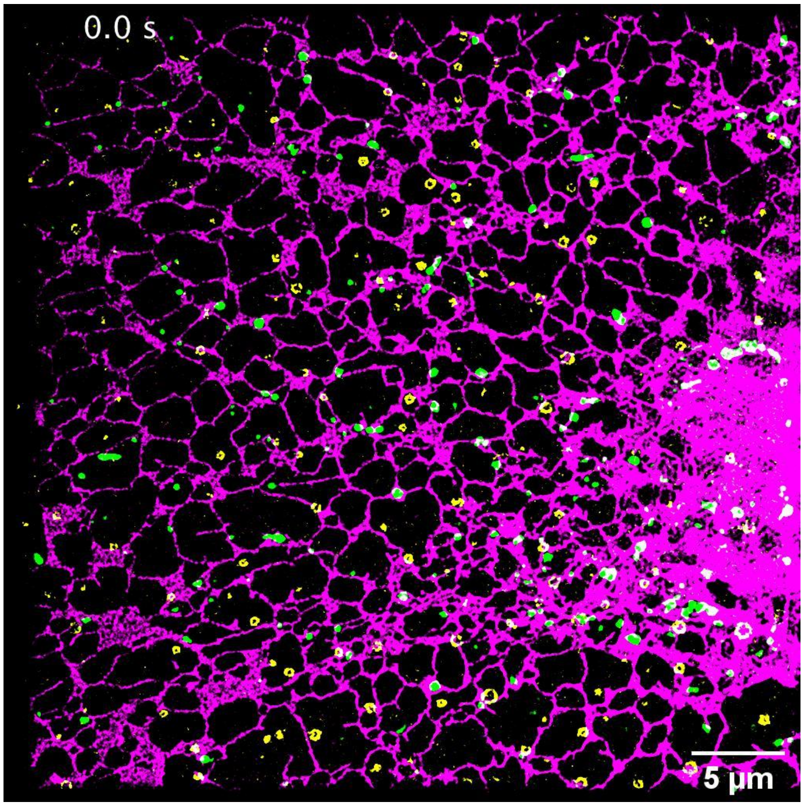


**Movie S2.**
**Live cell imaging by SIM demonstrates the dynamic motions of early endosomes and lysosomes.** A COS-7 cell expressing Rab5-EGFP (yellow) and mApple-Sec61b-C1 (magenta), stained with SiR-lysosome (green) was imaged over 90 s at 1.5 s/frame. Data as such cells were extracted by TrackMate to compare the organelle dynamics between early endosomes and lysosomes. (see **Fig. S1**)

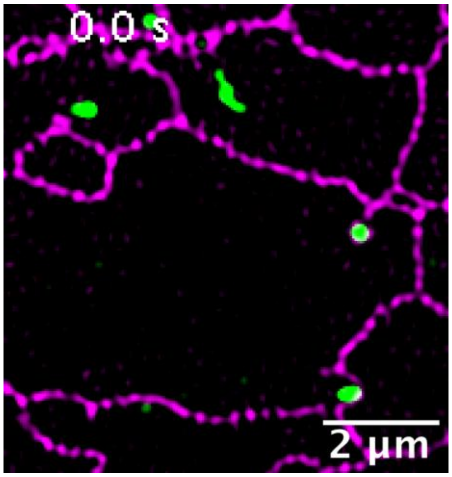


**Movie S3.**

**Live cell imaging by SIM demonstrates the coupled motions of the growing tip of a newly formed ER tubule (magenta) and associated lysosome (green).**

A region of a COS-7 cells expressing EGFP-hVAPA (magenta) and lysosomes labeled with SiR dye (green) was imaged over 90 s at 1.5 s/frame. The tip of ER forms three-way junctions. (see **Fig. 1B**)

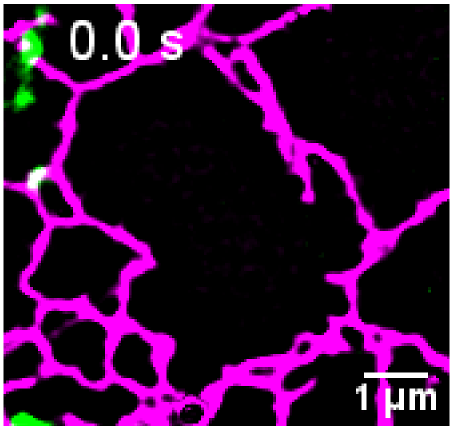


**Movie S4.**

**Live cell imaging by SIM demonstrates retraction of ER tubules when not associated lysosomes**.

A region of a COS-7 cells expressing EGFP-hVAPA (magenta) and lysosomes labeled with SiR dye (green) was imaged over 90 s at 1.5 s/frame. The tubules are unstable and retracts after a period of elongation. (see **Fig. 1C**)

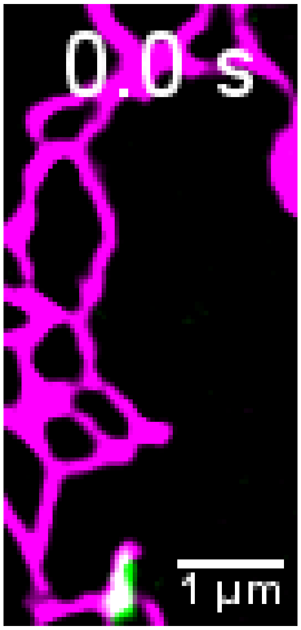


**Movie S5.**
**Live cell imaging by SIM demonstrates breakage of the connection between the growing tip of a newly-formed ER tubule and a lysosome.**
A region of a COS-7 cells expressing EGFP-hVAPA(KD/MD) (magenta) and lysosomes labeled with SiR dye (green) was imaged over 90 s at 1.5 s/frame. The tubule retracted after its detachment from a moving lysosome that was observed to gain an increase in velocity simultaneously. The lysosome positions at each time point were extracted by Trackmate to analyze the lysosome velocity changes (see **Fig. 2A**)

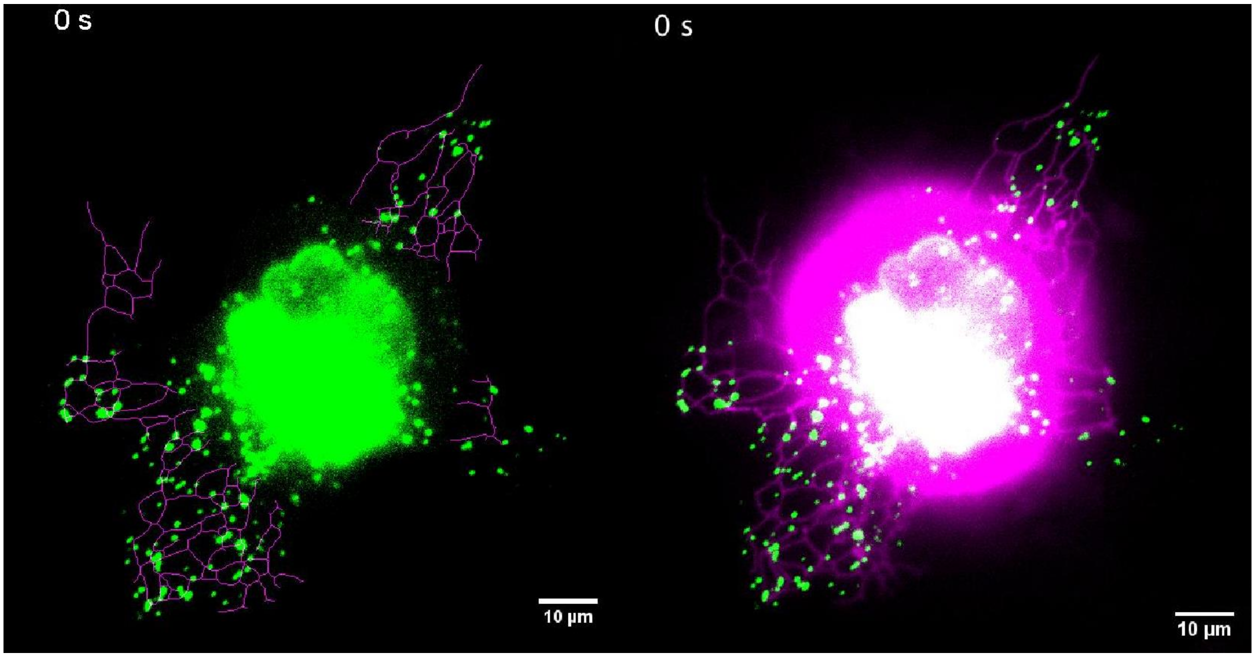


**Movie S6.**
**Live cell imaging of ontogenetically controlled lysosome motions and their guided ER reshaping.**
A COS-7 cell expressing LAMP1-mCherry-iLID (green), YFP-ER (magenta) and KIF1-VVDfast were imaged by widefield microscope over 4 mins at 2 s/frame. Left: ER channel of the widefield images was reconstructed to extract the skeletons by a trained artificial neural network for further quantification (see methods). Right: original widefield imaging.

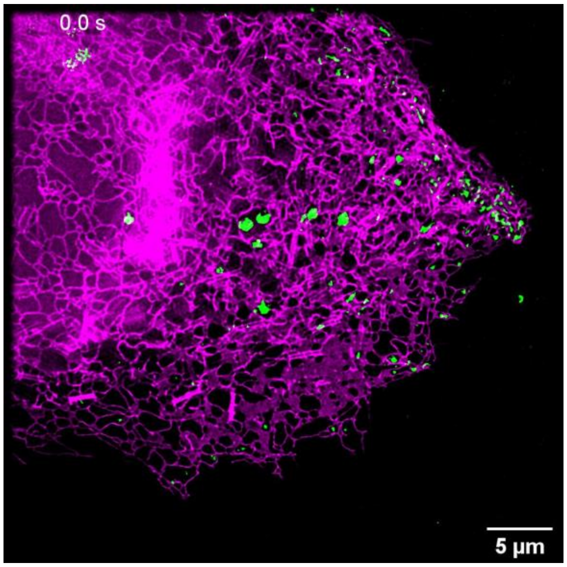


**Movie S7.**
**Live cell imaging by SIM demonstrates anterograde motions of lysosomes and ER network protrution under prolonged starvation.**
COS-7 cells expressing EGFP-hVAPA (magenta) and lysosomes labeled with SiR dye (green) were treated with serum-free DMEM for 24 hr then imaged over 90 s at 1.5 s/frame. The lysosome positions at each time point were extracted by Trackmate to analyze the movement direction (see **Fig. 3B3** and **S5**)

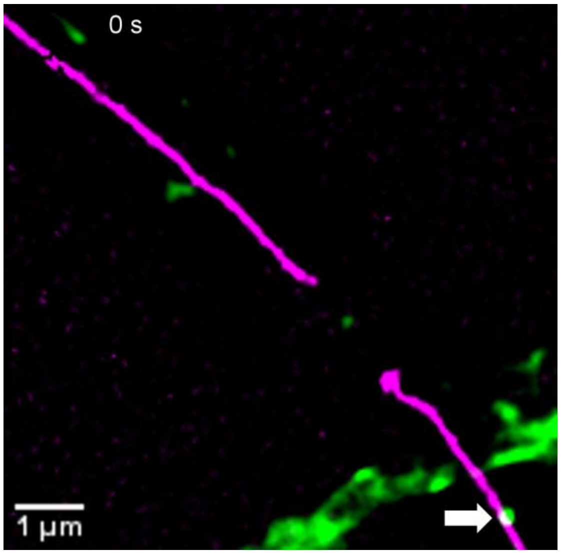


**Movie S8.**
**Live cell imaging by SIM demonstrates the reconnection of ER tubules by a moving lysosome in an VAPA expressing RGC axon.**
RGC axons expressing EGFP-hVAPA (magenta) and stained with SiR-Lysosome (green) were imaged at 5 s/frame. (see **Fig. 4Bi**)

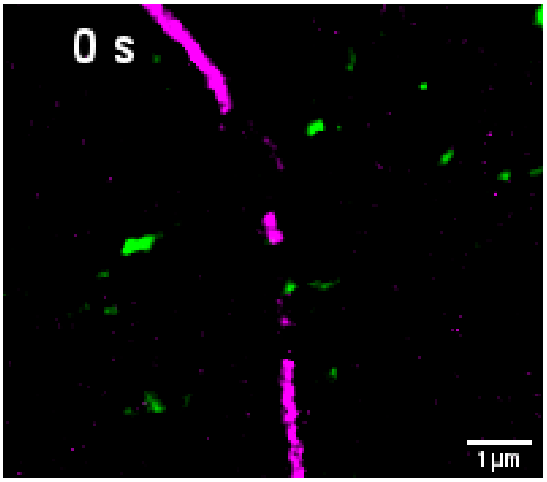


**Movie S9.**
**Live cell imaging by SIM demonstrates the persistent breakage of ER tubules in an VAPA KD/MD expressing RGC axon.**
RGC axons expressing EGFP-hVAPA KD/MD (magenta) and stained with SiR-Lysosome (green) were imaged at 5 s/frame. (see **Fig. 4Bii**)
